# Posterior specification of multi-lineage axial assembloids from human pluripotent stem cells

**DOI:** 10.1101/2024.08.29.610410

**Authors:** N Kee, M Leboeuf, S Gómez, C Petitpré, I Mei, S Benlefki, D Hagey, JM Dias, F Lallemend, S EL Andaloussi, J Ericson, E Hedlund

**Affiliations:** Department of Biochemistry and Biophysics, Stockholm University, Stockholm, Sweden; Department of Cellular and Molecular Biology, Karolinska Institutet, Stockholm, Sweden; Department of Neuroscience, Karolinska Institutet, Stockholm, Sweden; Department of Laboratory Medicine, Karolinska Institutet, Huddinge, Sweden; School of Biomedical Sciences, The University of Queensland, Australia

## Abstract

Elongation of the posterior body axis is driven by multi-potent neuromesodermal progenitors (NMPs), which both self-renew and simultaneously generate neural tube, neural crest, and presomitc mesoderm lineages at successive anterior posterior (A-P) levels. The ensuing diversification of these three NMP lineages is remarkably extensive, and also essential for an immense range of clinically important adult posterior body tissues. Here, we describe a human pluripotent stem cell protocol that successfully specifies authentic NMPs using a cocktail of seven factors (7F). 7F-NMPs express requisite markers, exhibit co-linear *HOX* activation, and can be purposely specified into each of the three NMP daughter lineages, demonstrating genuine multi-potency. 3D assembly of neural tube, neural crest, and presomitic mesoderm spheroids followed by long-term floating culture derives mature, multi-compartment Posterior Axial Assembloids, or PAXAs. PAXAs constitute a complex heterogeneous tissue containing spinal motor neurons and interneurons, central and peripheral glia, connective tissues, muscle satellite cells and contractile muscle fibres. Together, 7F-NMP and PAXA protocols establish a versatile in vitro platform to model mechanisms of human posterior body axis development, and for the study of a wide range of human diseases.

## Introduction

Modelling of early human development is encumbered by the ethical and technical limitations of accessing early primary tissue. In its place, human pluripotent stem cell (hPSC) based models have proven capable of recapitulating early embryonic development^1,2,3,4^. However, these models include gastrulation events, where the primitive streak breaks symmetry in the epiblast to form the cranial body axis. In vivo, this is followed by construction of the post-cranial axis, which is built up through the proliferative action of the caudal growth zone, a heterogeneous progenitor niche that emerges at the embryo’s most posterior point^5,6^. Major germ layers are represented in both cranial (anterior) and post-cranial (posterior) body axes, e.g., brain vs spinal cord^7^, stomach vs colon^8,9^, multiple neural crest derivatives^10^, or head and neck vs hindlimb musculature^11^. However, post-cranial lineages have taken a developmental path that is distinct from their cranial equivalents, namely through an intermediate caudal growth zone progenitor.

Here, we describe a set of human in vitro protocols that successfully and specifically recreates post-cranial events, through recapitulating the developmental lineage tree of one such caudal growth zone progenitor subtype: neuromesodermal progenitors (NMPs). NMPs are a small, transient and highly proliferative cell population that emerges after gastrulation and resides lateral to the regressing primitive streak in the anterior caudolateral epiblast^12,13^ (Fig. 1a). NMPs constitute a multipotent progenitor pool that self-renews and generates daughter progenitors of the neural tube, neural crest, and presomitic mesoderm lineages^14,13^. These daughter progenitors progressively integrate into the extending body axis, building up tissues in an anterior to posterior (A-P) fashion. Moreover, daughter progenitors specified at the same time express a matched *HOX* code that is instructive for their A-P identity^15^. Thus, all generated tissues vary in a coordinated fashion along the A-P axis, including e.g., spinal circuits, sympathetic ganglia, and bone and muscle structures. Recapitulating human NMP biology has therefore been the focus of many research efforts seeking to model post-cranial neural^16,17,18,19,20^, neural crest^21,22,23^, or neuromuscular tissue^24,25,26,27,28,29,30,31,32^.

**Figure 1.**
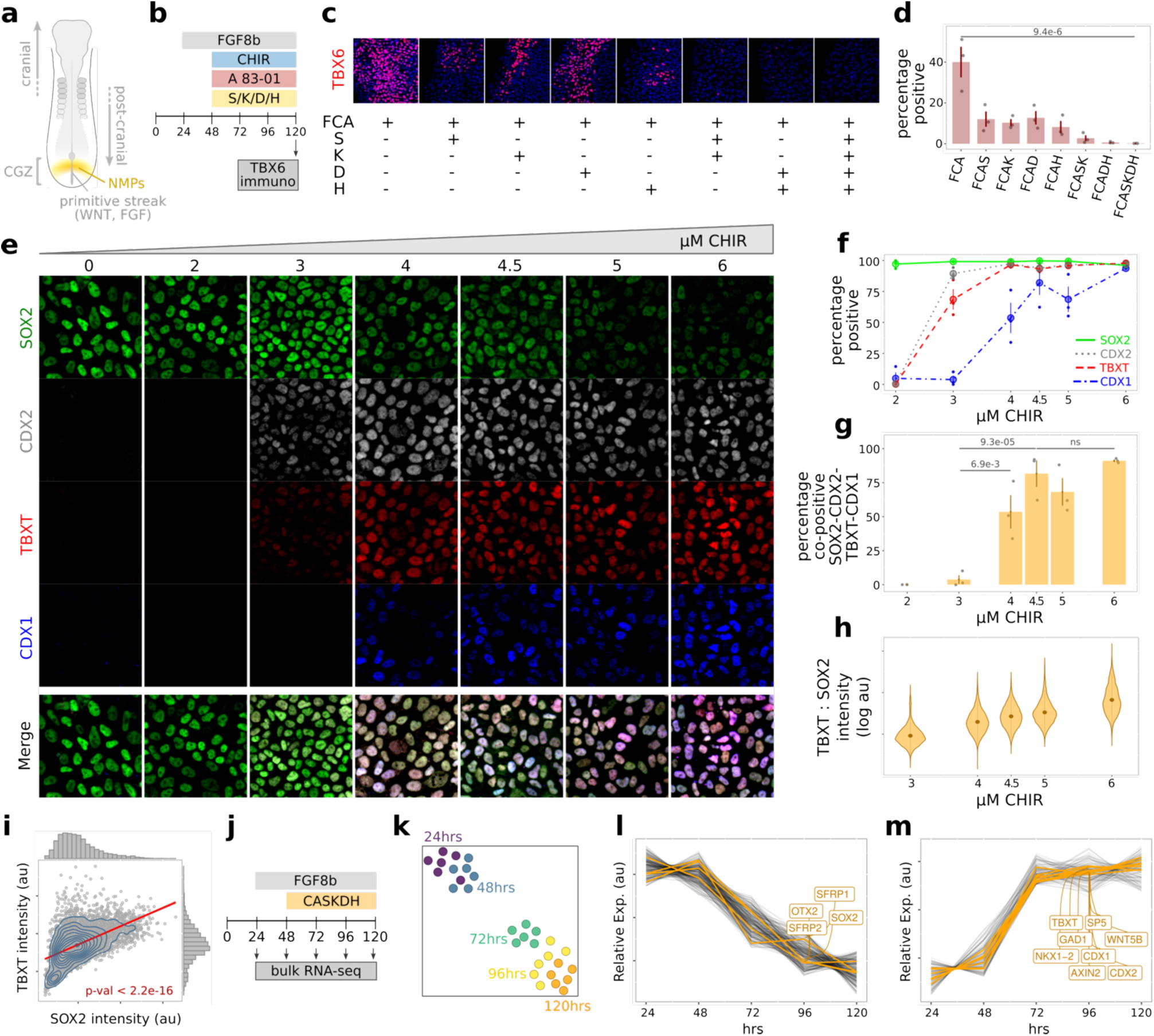
**(a)** Schematic of a developing human embryo showing NMPs located lateral to the regressing primitive streak. (**b)** Timing of factors used to specify NMPs using Fgf8b (F), CHIR (C), A 83-01 (A) and permutations of SIS3 (S), K02288 (K), DAPT (D) and heparin (H). (**c-d)** Combinatorial addition of components from 7F progressively eliminates off-target TBX6^+^ cells. **(e)** When applying this 7F cocktail, increasing concentrations of CHIR decrease the nuclear SOX2 intensity, while up-regulating expression of CDX2, TBXT and CDX1. **(f)** CDX2 and TBXT are robustly expressed at 3μM CHIR and above, while CDX1 is expressed beginning at 4μM CHIR and above. **(g)** SOX2-CDX2-TXBT-CDX1 co-positive cell frequency along the CHIR gradient. **(h)** In TBXT-SOX2 double positive cells, increasing concentrations of CHIR within 7F increases the TBXT:SOX2 intensity ratio. **(i)** At 4.5μM CHIR, both SOX2 and TBXT per-cell intensities positively correlate, and each display a normal distribution, indicating balanced expression. **(j)** Timing of factors used to pattern NMPs according to the optimised 7F cocktail, including time-points taken for bulk RNA-seq. **(k)** UMAP projection of bulk RNA-seq. Time-dependant gene expression analysis identifies gene sets that **(l)** decrease, or **(m)** increase, as NMP specification proceeds.

NMPs have previously been specified in vitro, from both mouse and human PSCs, using protocols incorporating CHIR99021 (CHIR), a WNT pathway activator, and FGFs^24,16,25^. This combination induces expression of CDX2, which together with CDX1 is crucial for in vivo co-linear *HOX* activation and post-cranial specification^33^. CHIR and FGF also induce co-expression of the neural determinant SOX2 and the mesendodermal determinant TBXT. These two transcription factors co-occupy many NMP expressed genes^34,35^, are sustained by TBXT/CDX/WNT/FGF autoregulatory feedback loops^36,37,34^, and mutually antagonise neural vs mesodermal specification^38,34,39,35^. As such, balanced SOX2/TBXT co-expression in NMPs is a hallmark of multi-potency. Studies have also shown that, when added in addition to CHIR and FGF, both dual SMAD inhibition^28^ and Notch inhibition^32^ suppress TBXT, biasing NMPs towards a neural program. In further reports, neural induction of NMPs can be achieved through addition of retinoic acid^40^ (which abrogates FGF and WNT signalling in vivo to dismantle the NMP niche^41,42^), allowing derivation of neural tube progenitors amenable to BMP mediated dorsalisation, and/or sonic hedgehog mediated ventralisation^24,16,25,43,44,45^. NMP-like cells can also be specified into the neural crest lineage through application of BMP ligands^12,22,23^, while sustained CHIR exposure has been used in multiple protocols to specify presomitic mesoderm^46,47,48^, at times passing through a short, transitory NMP-like state^49^.

However, while all arms of the NMP lineage tree have been recapitulated, this requires multiple protocols that often use different base media and coatings, or that specify downstream lineages in different mixtures and/or with different timings. Moreover, there are no reports of an efficient protocol that can derive all lineages from the same starting NMP. This discordance has greatly hindered the study and use of posterior human tissues and raises concerns regarding the authenticity of these protocols’ NMP-like cells. Further, modelling of different protocols’ in vitro NMP state suggests it constitutes an unstable shallow attractor^50^, that is subtly sensitive to secreted WNT and FGF ligands^51^. In line with this, in vitro posterior human tissues often contain cells with a mix of A-P levels^18,30,31^, suggesting that not all NMPs are stably maintained and instead spontaneously differentiate into downstream lineages. Here, we overcome these obstacles by identifying unified culture conditions that recapitulate the hallmarks of authentic NMPs: stable maintenance of NMP state, temporal co-linear *HOX* induction, and genuine multipotency allowing efficient and controlled specification into post-cranial neural tube, neural crest and presomitic mesoderm lineages.

## Results

### In vitro specification of human NMPs using a cocktail of seven factors

To identify in vitro culture conditions that specify authentic NMPs, we differentiated hPSCs^52^ using a CHIR pulse beginning at 48hrs^16^, together with combinatorial screening of additional small molecules (Fig. 1b). Our baseline cocktail contained three factors: Fgf8b, CHIR and A 83-01 (an inhibitor of the ACTIVIN/NODAL/TGFβ pathway) (FCA). These conditions gave NMP-like cells co-expressing SOX2 and TBXT, however some cells displayed higher levels of either SOX2 (filled arrowhead) or TBXT (un-filled arrowhead) (Extended Data Fig. 1a-b). Moreover, many cells located towards the edge of the colonies expressed high levels of TBXT, no SOX2, and instead expressed the presomitic mesoderm determinant TBX6 (arrow), indicating they had left the NMP state. We therefore screened additional small molecules that would eliminate off-target TBX6+ cells (Fig. 1c-d, Extended Data Fig. 1c), and identified four additional factors: SIS3 (S), an inhibitor of intracellular SMAD3; K02288 (K), an inhibitor of cell surface ALK1/2/6 receptors; DAPT (D), an inhibitor of the NOTCH effector, γ-secretase; and heparin (H), a large and complex molecule enriched in the extracellular matrix. In FCA conditions ∼40% of cells expressed TBX6. Addition of either S, K, D or H reduced this to ∼10%. Combinatorial addition of 5F (FCASK or FCADH) improved NMP purity even further, while the combined 7F cocktail FCASKDH effectively eliminated all TBX6+ cells (Fig. 1d).

### Graded CHIR concentrations affects 7F-NMP marker expression

When applying 7F, we also discovered that the concentration of CHIR greatly affected the expression of NMP marker genes (Fig. 1e-f). Automated analysis of SOX2, CDX2, TBXT, and CDX1 immunostainings showed that while SOX2^+^ cells were detected independent of CHIR concentration, CDX2^+^ and TBXT^+^ cells were first detected at 3µM CHIR (a concentration of CHIR routinely used for in vitro monolayer NMP induction). In contrast, CDX1 was first detected at a higher concentration of 4µM CHIR, where it marked ∼50% of cells. This increased to ∼80% of cells at 4.5µM CHIR, and ∼90% at 6µM CHIR. Further, between 4-6µM CHIR all CDX1^+^ cells were also SOX2^+^/TBXT^+^/CDX2^+^ (Fig. 1g). CDX1 induction only at higher CHIR levels led us to compare the expression patterns of *SOX2*, *CDX2* and *CDX1* together in vivo (Extended Data Fig. 1d). RNA scope analysis in E8.5 mouse caudal growth zone tissue showed that *SOX2* and *CDX2* mRNA were broadly co-expressed in all caudal growth zone cells (closed arrowheads), while *CDX1* mRNA was enriched in NMPs in the pseudo-stratified surface caudolateral epiblast (open arrowhead), in line with previous reports^53^. Thus, co-expression of CDX1 together with SOX2/TBXT/CDX2 constitutes a stringent NMP marker profile. Next, we tested different FGF receptor ligands, since there is no consensus in NMP protocols regarding the use of FGF2 vs FGF8b. We found that substitution of FGF8b (a caudal growth zone growth factor) for FGF2 led to re-emergence of TBX6^+^ cells (Extended Data Fig. 1e), again at the periphery of the colony, and thus FGF8b outperforms FGF2 in preventing off-target TBX6+ presomitic mesoderm. Further, we found that as CHIR concentration increased between 3 to 6µM, the per cell expression level of SOX2 decreased, while the per cell TBXT expression level increased (Extended Data Fig. 1g-h), effectively leading to an increase in the TBXT:SOX2 ratio (Fig. 1h). Lastly, these CHIR dependent responses of SOX2, TBXT, CDX2, and CDX1 expression were all recapitulated in a second hPSC line^54^ (Extended Data Fig. 1f).

We next sought to choose a CHIR concentration that would not inappropriately bias the multi-potency of our 7F-NMPs. We decided upon 4.5µM CHIR, since this captured an intermediate expression level along the CHIR concentration gradient for both TBXT and SOX2 (Extended Data Fig. 1g-h). Importantly, at 4.5µM CHIR, the per-cell expression levels of SOX2 and TBXT both showed a normal distribution that was positively correlated (p= 2.2e-16, Fig. 1i), indicating that at the single-cell level SOX2 and TBXT expression were balanced. When considering the entire gradient, 4.5µM CHIR showed a linear relationship between TBXT and SOX2 levels that could be down-shifted (lower TBXT) by decreasing CHIR or left-shifted (lower SOX2) by increasing CHIR (Extended Data Fig. 1i-i’), suggesting that 7F-NMPs can occupy a range of neural-to-mesodermal bias depending on levels of WNT signalling. Nonetheless, these observations indicated that at 4.5µM CHIR, TBXT and SOX2 were stably co-expressed within individual 7F-NMPs.

### Bulk RNA-seq of 7F-NMPs confirms upregulation of requisite markers

Next, we investigated the transcriptome-wide RNA profile of 7F-NMPs through their full differentiation time course. Cells were collected at 24hr timepoints between 24-120hrs (Fig. 1j) and processed as mini-bulk samples using Smart-seq3 single-cell RNA-seq (scRNA-seq) chemistry^55^. UMAP analysis projected samples in a linear fashion according to their timepoint (Fig. 1k), with pre 7F timepoints (24 and 48hrs) in one corner, and post 7F timepoints (72, 96, 120hrs) in the opposite corner. Time dependant expression analysis identified gene sets indicative of this transition. In a gene set that was downregulated post 7F (Fig. 1l), we noted the primed epiblast marker *OTX2*, and the secreted WNT inhibitors *SFRP1/2*, which are expressed in the anterior cranial neurectoderm.

Also downregulated was *SOX2*, which displays distinct chromatin occupancy depending on its expression level^35,56^. At high levels SOX2 maintains pluripotency, while reduced levels change its occupancy to caudal epiblast expressed genes, where it co-occupies low-affinity SOX binding sites alongside TBXT and CDX. Next, we identified a gene set that was upregulated after addition of 7F and that contained multiple NMP markers (Fig. 1m). As expected, both *CDX1* and *CDX2* were upregulated, indicating post-cranial specification. Upregulation of *WNT5B* and its target gene *TBXT* are consistent with expansion of the NMP pool^5^, while *SP5* activates WNT target genes^57^. *GAD1* is a direct target of b-catenin^58^ and has previously been identified as a marker of human NMPs^59^. *NKX1-2* was also in this gene set, and genetic lineage tracing in mice demonstrates that NKX1.2^+^ cells contribute to all A-P levels of the post-cranial axis^60^. Lastly, this list overlaps considerably with previous reported NMP marker profiles from mouse^49^ and chicken^13^ (20 genes occurring in at least two of the three gene lists, including *TBXT, CDX2*, *NKX1-2*, *WNT5B*, *SP5*, *AXIN2*, and *GAD1*), corroborating our cultures’ differentiation into an NMP identity.

A further trait of NMPs is their capacity to undergo temporal co-linear *HOX* activation while maintaining their multipotency. This enables continuous derivation of neural tube, neural crest and somitic mesoderm with distinct A-P identities. In embryos, *Hox* expression is initiated by Wnt signalling in the posterior epiblast^61,62^ and is required for NMP maintenance and axial elongation^63,37^. As elongation begins, expression of 3’ Hox genes mark anterior structures, while later in time their 5’ neighbours are upregulated, leading to a nested *Hox* code that specifies progressively more posterior structures. CDXs, while dispensable for initial opening of 3’ regions of the A and B *Hox* clusters, are required for DNA accessibility within middle regions, where they propagate co-linear activation^64^. Further, lumbo-sacral *Hox* induction across the trunk to tail transition is controlled by GDF11^65^, a TGF-β member upregulated in the caudal growth zone beginning at sacral stages^66^. That *CDX1/2* were expressed in our cultures suggested that co-linear *HOX* activation may be occurring in 7F-NMPs. We therefore investigated the progression of HOX gene expression in our 7F-NMP RNA-seq data, and also in NMPs differentiated for a further 48hrs in a six factor cocktail (6F) that omitted two inhibitors of TGF-β signalling (A 83-01 and SIS3) while adding GDF11. Analysis of 7F-to-6F NMP RNA-seq data showed upregulation of *HOX1-2* genes within 24hrs of 7F exposure. *HOX3-9* genes were sequentially upregulated over the next 48hrs, while *HOX10* transcripts were only detected after addition of 6F (Extended Data Fig. 1j).

In total, immunostainings of requisite markers and transcriptome-wide bulk RNA-seq analysis corroborates a high efficiency induction of 7F-NMPs that hold key traits of their in vivo counterparts. Namely, expression of cross-species NMP markers at both the protein and RNA levels, while also exhibiting temporal co-linear *HOX* activation consistent with A-P specification from cervical to lumbar (with 6F) levels.

### Specification of 7F-NMPs into presomitic mesoderm requires WNT, FGF, and extracellular matrix factors

Authentic in vitro NMPs should be able to efficiently differentiate into all downstream lineages expected from the vertebrate embryo (Fig. 2a). We began by testing 7F-NMPs’ potential to generate presomitic mesoderm by culturing cells using different factors between 120-192hrs, and screening for TBX6^+^ cells (Fig. 2b’-b’’, Extended Data Fig. 2a-c). We investigated three factors previously implicated in *in vitro* differentiation of presomitic mesoderm; CHIR^47^, FGF2^46^, and basal matrix extract (BME), a complex mix of extracellular matrix molecules which have previously been shown to affect the morphogenesis of 3D cultured post-cranial structures^67,30,31^, and to promote epithelial-mesenchymal transition (EMT) behaviours^68^. Maintenance of 7F-NMPs in base media alone did not induce TBX6 expression. Activating WNT signalling alone using a high concentration of 10µM CHIR induced only a modest increasing trend to ∼12% TBX6^+^ cells. Similarly, addition of FGF2, or drop application of a gelled layer of BME alone, had no impact on TBX6 induction. However, the combined action of all three (CFB) increased the number of TBX6^+^ cells to ∼80%. Further, neither CF, CB, nor FB could induce increased levels of TBX6 when compared to CFB (Extended Data Fig. 2b-c). We further found that the concentration of CHIR in the CFB cocktail was crucial, since decreasing CHIR from 10 to 4.5µM reduced the proportion of TBX6^+^ cells to ∼25%. Similarly, substitution of FGF2 for FGF8b reduced the proportion of TBX6^+^ cells to ∼10%, while substitution of BME for Collagen I reduced the proportion of TBX6^+^ cells to ∼20%. Inhibition of SMAD signalling has been previously reported in presomitic mesoderm protocols^46,47,48^, however we found that addition of A83-01 or K02288 on top of CFB did not affect TBX6 induction (Extended Data Fig. 2b-b’). Importantly, we also found that CFB induction of TBX6 was dependant on the concentration of CHIR used during the prior 7F-NMP phase (Fig. 2b’’, Extended Data Fig. 2a). That is, if 7F-NMPs were generated using a lower concentration of 3µM CHIR, TBX6 induction by subsequent CFB reduced to ∼30%. Thus, CFB (applied to 7F-NMPs derived using 4.5µM CHIR) induced the highest proportion of TBX6+ cells, and also displayed the highest per cell intensity of TBX6 (Extended Data Fig. 2a’, 2c’).

**Figure 2.**
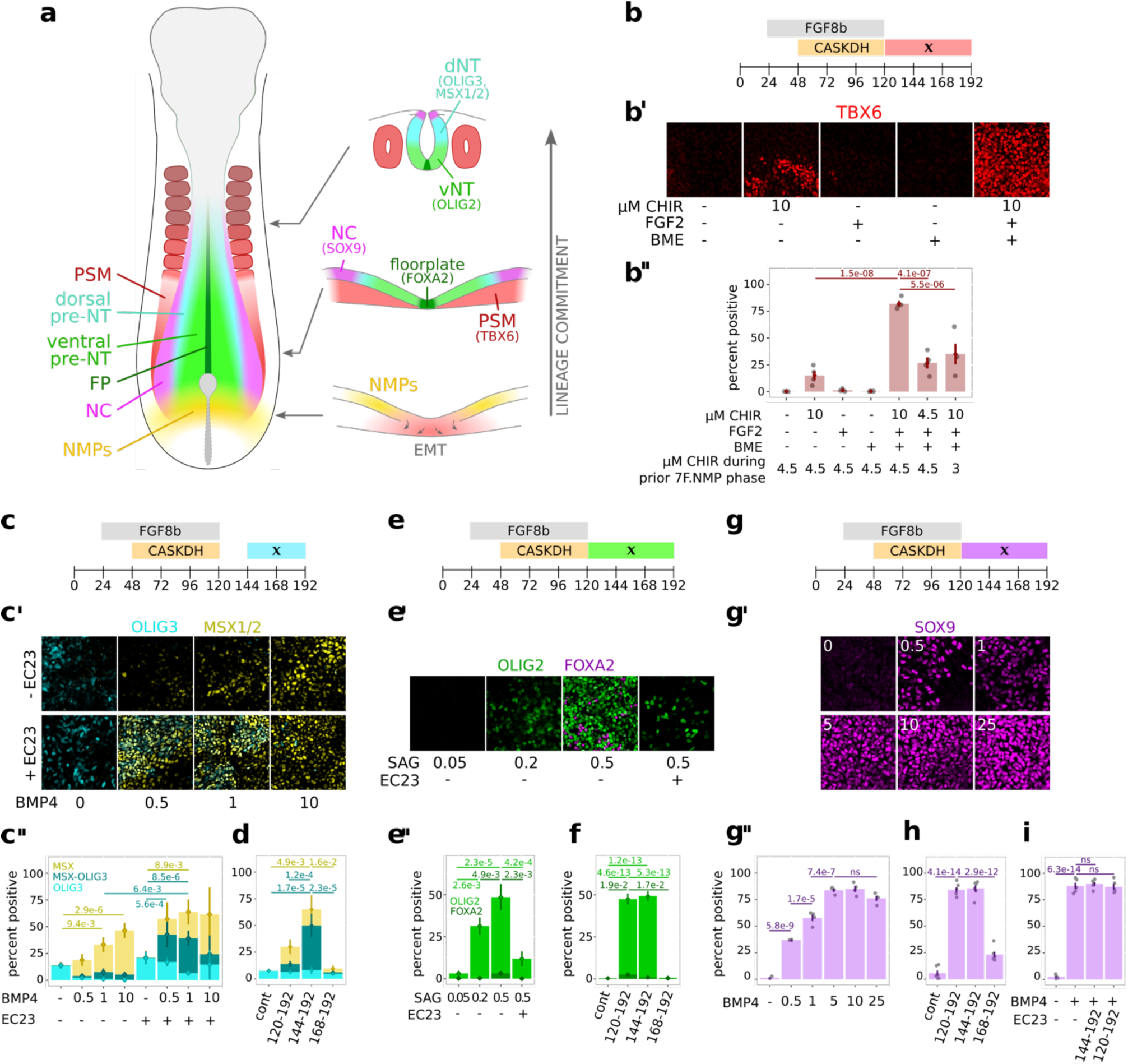
**(a)** Developing human embryo schematic depicting NMPs in the surface caudolateral epiblast of the caudal growth zone, and their daughter presomitic mesoderm (TBX6), neural crest (SOX9), floor plate (FOXA2), ventral neural tube (e.g., OLIG2), and dorsal neural tube (e.g., OLIG3) lineages. **(b)** Timing of factors used for presomitic mesoderm differentiation. *x* denotes the period where different permutations of CHIR, FGF2, or Basal Matrix Extract (BME), are applied. **(b’)** TBX6 immunostaining showing presomitic mesoderm progenitors emerging in the different permutations. **(b’’)** Quantification of b’, showing the highest proportions of TBX6^+^ cells with combined CFB. Notably, maximal presomitic mesoderm induction requires 4.5μM CHIR in the prior 7F-NMP phase. **(c)** Timing of factors used for dorsal neural tube differentiation. *x* denotes the period where different permutations of BMP4 (ng/ml) and EC23 are applied. **(c’)** OLIG3/MSX1/2 immunostaining showing dorsal neural tube progenitors emerging in different permutations. **(c’’)** Quantification of OLIG3/MSX1/2^+^ cells in c’, showing the highest proportions with BMP4 (0.5-1ng/ml) + EC23. **(d)** Differentiating dorsal neural tube generates the highest proportions of OLIG3/MSX1/2^+^ cells when BMP4(1ng/ml) + EC23 is applied between 144-192hrs. **(e)** Timing of factors used for ventral neural tube differentiation. *x* denotes the period where different permutations of SAG(μM) or EC23 are applied. **(e’)** OLIG2/FOXA2 immunostaining showing the emergence of ventral neural tube progenitors in different permutations. **(e’’)** Quantification of OLIG2^+^ and FOXA2^+^ cells in e’, showing the highest proportions with 0.5μM SAG, which is abolished by addition of EC23. **(f)** Differentiating ventral neural tube generates the highest proportions of OLIG2^+^ and FOXA2^+^ cells when 0.5μM SAG is applied between 120-192hrs. **(g)** Timing of factors used for neural crest differentiation. *x* denotes the period where different permutations of BMP4 and/or EC23 are applied. **(g’-g’’)** BMP4 increases the proportion of SOX9^+^ neural crest progenitors in a dose dependant manner. **(h)** Robust SOX9^+^ neural crest induction requires early addition of BMP4 (10ng/ml) between 120/144hrs-192hrs. **(i)** Addition of EC23 immediate after exit from 7F-NMP (120-192hrs), or after a 24hr delay (144-192hrs), does not affect the efficiency of neural crest induction.

### Specification of 7F-NMPs into dorsal neural tube requires delayed BMP and retinoic acid signalling

We next investigated the potential of 7F-NMPs to generate dorso-ventralised neural tube. In vivo, neural tube patterning occurs in two phases^69^. The early phase is termed the pre-neural tube, where cells that have only recently exited the NMP niche are ventralised by sonic hedgehog released from the notochord^70,71^. The later phase occurs in the neural tube proper where sonic hedgehog released from the floor plate continues to pattern ventral neural tube progenitors^70, 71^, while BMP released from the roof plate patterns dorsal neural tube progenitors^72^, and retinoic acid is also being released from adjacent somites^41^. Thus, we began patterning dorsal neural tube progenitors by recapitulating this later phase. We tested the dorsalising factor BMP4 and the highly stable retinoic acid receptor agonist EC23 between 144-192hrs (Fig. 2c-c’’, Extended Data Fig. 2d). As a readout, we assessed the induction of OLIG3, which marks d1-3 progenitor domains^73,74^, and MSX (using an antibody against both MSX1 and MSX2) which marks d1-4 progenitor domains and the roof plate at various stages of development^75^. Maintaining 7F-NMPs in base media induced ∼10% OLIG3^+^ cells but did not induce MSX1/2. Adding BMP4 alone did not change the proportions of OLIG3^+^ cells at any concentration but did induce MSX1/2 in a dose dependant manner from ∼15% at 0.5 ng/ml, to ∼30% at 1 ng/ml, and ∼50% at 10 ng/ml. EC23 alone did not affect the proportion of OLIG3^+^ cells compared to base media, and also failed to induce expression of MSX1/2. However, double labelled cells were observed when combining both BMP4 and EC23. Here, BMP4 induced OLIG3^+^/MSX1/2^+^ cells to ∼25% at 0.5ng/ml, and ∼35% at 1ng/ml. Next, we investigated the time window within which 7F-NMPs were sensitive to combined BMP4 (1ng/ml) and EC23 (BE) and found that double labelled OLIG3^+^/MSX1/2^+^ cells were diminished when BE was applied immediately after 7F-NMP conditions (120-192hrs), and also 48hrs after leaving them (168-192hrs) (Fig. 2d). Instead, maximal induction was achieved only when BE was applied 24hrs after 7F-NMP exit (144-192hr), recapitulating the later phase of dorsal neural tube patterning observed *in vivo*.

### Specification of 7F-NMPs into ventral neural tube requires early sonic hedgehog signalling

Next, we sought to recapitulate the early phase of pre-neural tube ventral patterning by applying the sonic hedgehog pathway activator SAG, immediately after NMP exit between 120-192hrs (Fig. 2e-f, Extended Data Fig. 2e-f). As readout, we screened for the motor neuron progenitor marker OLIG2, and the floor plate marker FOXA2. A low dose of 50nM SAG resulted in only ∼2% OLIG2^+^ cells, while increasing this to 200nM generated ∼30% OLIG2^+^ cells, and 500nM generated ∼50% OLIG2^+^ cells. FOXA2 positive floor plate progenitors were also maximally induced at the highest 500nM dose. Moreover, 500nM SAG induced the highest per cell intensities for both OLIG2 and FOXA2 (Extended Data Fig. 2e’-e’’). In vivo, induction of the floor plate in the early pre-neural tube requires FGF signalling^69^, which is abrogated by retinoic acid released from anterior somites^76^. Further, retinoic acid released at neural tube levels constrains expression of Shh in the ventral-most neural tube^77^. In line with these observations, addition of EC23 immediately after exit from NMP conditions eliminated almost all FOXA2^+^ floor plate progenitors, and significantly decreased the number and staining intensities of OLIG2^+^ motor neuron progenitors (Fig. 2e, Extended Data Fig. 2e). Next, we investigated the time window within which 7F-NMPs were sensitive to sonic hedgehog signalling. In line with early pre-neural tube ventralisation *in vivo*, cells were amenable to OLIG2 induction if SAG was given early, either between 120-192hrs or 144-192hrs (Fig. 2f, Extended Data Fig. 2f). FOXA2 induction was maximal at 120-192hrs, while SAG addition at 168-192hrs generated effectively no OLIG2 or FOXA2 cells. Thus, ventral patterning of 7F-NMPs using SAG between 120-192hrs recapitulates the early phase of pre-neural tube patterning observed *in vivo*.

### Specification of 7F-NMPs into neural crest requires early BMP signalling

Finally, we directed the differentiation of 7F-NMPs towards the neural crest lineage. In vivo, neural crest is specified in the early pre-neural tube, through activation of the BMP pathway^72^. We added graded concentrations of BMP4 immediately after NMP exit between 120-192hrs and found that SOX9^+^ cells were induced in a dose dependant manner (Fig. 2g-g’’, Extended Data Fig. 2g). Neural crest induction peaked at 5ng/ml, where ∼85% of the cells were SOX9^+^. In line with in vivo expectations, addition of BMP4 (10ng/ml) between 120-192hrs or 144-192hrs again induced ∼85% SOX9^+^ cells, but this competence was lost at 168hrs (Fig. 2h). Furthermore, while no off-target OLIG3^+^ dorsal neural tube progenitors were observed when BMP4 was applied between 120-192hrs, occasional OLIG3^+^ cells emerged when BMP4 application was delayed (Extended Data Fig. 2h). Lastly, retinoic acid has also been shown to be crucial for maturation of neural crest progenitors^78^, and in line with this addition of EC23 had no influence on SOX9 induction (Fig. 2i).

In total, these four monolayer protocols demonstrate that 7F-NMPs specified using 4.5µM CHIR can be differentiated in a controlled manner into all downstream NMP lineages. Further, this specification recapitulates temporal and dose dependant signalling paradigms observed in vivo and demonstrates that 7F-NMPs hold the key hallmark of multipotency.

### scRNA-seq interrogation of 7F-NMPs and their downstream lineages

Assessment of hPSC differentiation efficacy using immunohistochemistry can only evaluate markers that are intentionally selected. By contrast, RNA-seq allows assessment of marker genes transcriptome-wide. To interrogate our protocols, we applied Smart-seq3 to 7F-NMPs at 48, 72, 96 and 120hrs of culture, and to presomitic mesoderm, ventral neural tube, dorsal neural tube and neural crest at 144, 168 and 192hrs of culture (Fig. 3a). UMAP analysis and graph-based Infomap clustering of all 6022 cells showed distinct clusters of NMPs (NMP.1-3), presomitic mesoderm (PSM.1-3), two pre-neural tube clusters (preNT.1A-B), ventral neural tube progenitors (vNT.2-3), dorsal neural tube progenitors (dNT.2-3), a small cluster of differentiating neurons (neurons), and a cluster of preNMPs (Fig. 3b). Dot plotting of all cell clusters showed that requisite markers for each cell type were expressed (Fig. 3c). We then investigated the purity of each of our differentiation protocols, finding that all protocols derived at least 95% of on-target cells at 192hrs, except for the dorsal neural tube protocol, which also included ∼45% off-target neural crest cells (Fig. 3d, Extended Data Fig. 3a).

**Figure 3.**
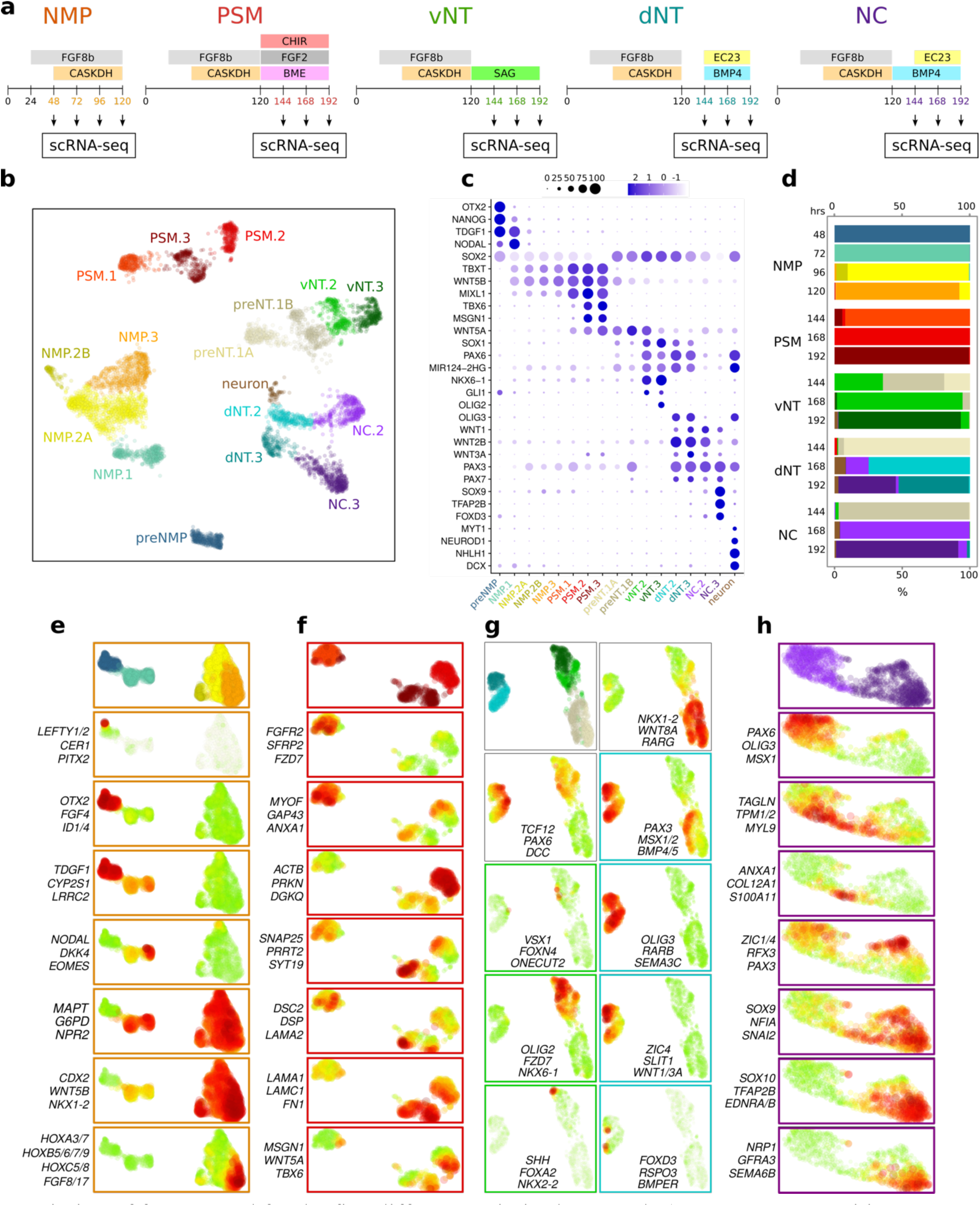
**(a)** Timing of factors used for the five different optimised protocols (7F-NMPs, presomitic mesoderm, ventral neural tube, dorsal neural tube and neural crest), including the time-points from which single cells were processed for Smartseq3 scRNA-seq. **(b)** Combined UMAP of cells from all five protocols. Infomap analysis identified 17 clusters (5x NMP, 3x presomitic mesoderm, 2x pre-neural tube, 2x ventral neural tube, 2x dorsal neural tube, 2x neural crest and 1x neurons). **(c)** Expression of requisite lineage markers across all 17 clusters. **(d)** Percentage contribution of each of the 17 clusters to each differentiation timepoint illustrates that all protocols, with exception of the dorsal neural tube protocol, has high on-target differentiation efficiency. Gene sets containing co-expressed genes were determined through sub-clustering of **(e)** only NMP clusters, **(f)** only presomitic mesoderm clusters, **(g)** combined pre-neural tube, ventral-neural tube and dorsal-neural tube clusters, and **(h)** neural crest clusters.

Next, we sub-clustered NMP cells, presomitic mesoderm cells, combined pre-neural tube, ventral neural tube and dorsal neural tube cells, and neural crest cells (Fig. 3e-h). Regarding NMPs (Fig. 3e), the preNMP cluster (majority 48hr cells) comprised cells exposed to only FGF8b for 24hrs and expressed gene sets containing multiple epiblast and pluripotency markers (*OTX2*, *FGF4*, *ID1/4*, *TDGF1*, *LRRC2*), as well as a gene set marking a small number of FGF8 dependant node genes involved in left-symmetry (*LEFTY1/2*, *CER*, *PITX2*). The NMP.1 cluster (majority 72hr cells) comprised cells exposed to only 24hrs of 7F, and expressed a gene set containing posterior epiblast markers such as *NODAL*, *DKK4*, and *EOMES*, suggesting a transition towards a caudal growth zone identity. NMP.1 .2 and .3 clusters comprised cells exposed to 7F for 24, 48 or 72hrs respectively, (majority 72hr, 96hr and 120hr cells) and all expressed a gene set containing cytoskeletal proteins and cell growth genes (*MAPT*, *G6PD*, *NPR2*). In contrast, only NMP.2 and .3 clusters expressed a gene set containing hallmark NMP markers such as *CDX2*, *WNT5B*, and *NKX1-2*, indicating that NMP specification requires exposure to 7F for at least 48hrs. There were no gene sets that obviously described lineage differences between NMP.2A and NMP.2B, with both groups showing expression of requisite NMP markers (Extended Data Fig. 3b). Differential gene expression and subsequent cellular component GO analysis identified NMP.2B enriched terms associated with focal adhesion and cell-substrate junctions (Extended Data Fig. 3c-c’’), however NMP.2B cells may also cluster apart due to a reduced number of detected genes (Extended Data Fig. 3c’’’). Lastly, the NMP.3 cluster (majority 120hr cells) expressed a gene set containing many post-cranial marker genes, including FGF8 and post-cranial HOX genes ranging up to *HOX9* thoracic A-P levels (Extended Data Fig. 3d-e).

Sub-clustering of presomitic mesoderm progenitors (Fig. 3f) identified a series of gene sets enriched at successive days of differentiation. Upregulated in PSM.1 cells was a gene set containing members of the FGF and WNT pathways (*FGFR2*, *SFRP2*, *FZD7*), and a gene set containing genes implicated in cell migration and EMT (*MYOF*, *GAP43*, *ANXA1*). Gene sets containing mediators of vesicle biology and exocytosis were found in either PSM.2 (*ACTB*, *PRKN*, *DGKQ*) or both PSM.2 and PSM.3 cells (*SNAP25*, *PRRT2*, *SYT19*). Further gene sets were enriched in PSM.3 cells that contained mediators of cell-cell adhesion (*DSC2*, *DSP*, *LAMA2*) or genes encoding extracellular matrix interacting proteins (*LAMA1*, *LAMC1*, *FN1*). Lastly, a gene set containing crucial markers of presomitic mesoderm identity (*MSGN1*, *WNT5A*, *TBX6*) was upregulated in cells of both PSM.2 and PSM.3. Together these results suggest that our cultures undergo an EMT, a process understood to involve dynamic cytoskeletal remodelling, secretion of extracellular matrix modifiers, and cell migration^79^, finally leading to upregulation of core transcriptional determinants of presomitic mesoderm.

Sub-clustering of combined pre-neural tube, ventral neural tube, and dorsal neural tube progenitors (Fig. 3g) identified multiple gene sets indicative of neural specification. preNT.1A and preNT.1B cells both expressed a gene set containing the preNT markers *NKX1-2* and *WNT8A*, while preNT.1B additionally expressed a number of BMP target genes (*PAX3*, *MSX1/2*, *BMP4/5*), in line with the majority of these cells belonging to the 144hr neural crest timepoint (Extended Data Fig. 3a). Both dNT.2-3 and vNT.2-3 expressed a gene set containing the pan-neural progenitor markers *TCF12*, *PAX6* and *DCC*, while dNT.2-3 also expressed gene sets for dorsal neural tube progenitors (*OLIG3*, *SEMA3A*) and the differentiating roof plate (*WNT1/3A*, *SLIT1*, *FOXD3*). vNT.2-3 clusters included cells expressing gene sets for ventral p2A markers (*VSX1*, *FOXN4*) and pMN markers (*OLIG2*, *NKX6.1*), while vNT.3 contained a small number of cells expressing a gene set for floor plate cells (*SHH*, *FOXA2*, *NKX2.2*). Thus, despite using a simplistic strategy for dorso-ventral specification, that is, a single timepoint for application of a single dose of dorsal (BMP4 + EC23) or ventral (SAG) patterning signals, multiple dorso-ventral progenitor subtypes were represented in our cultures.

Sub-clustering of all neural crest clusters (Fig. 3h) mapped NC.2 cells (majority 168hr) to one half of the UMAP, and NC.3 cells (majority 192hr) to the opposite half. In the centre of the UMAP was a small number of cells co-expressing markers for neural crest derived smooth muscle (e.g., *TAGLN*, *TPM1/2*, *ANXA1*, *COL12A1*). NC.2 cells expressed a gene set containing dorsal neural tube markers (*PAX6*, *OLIG3*, *MSX1*), while cells in both NC.2 and NC.3 expressed a gene set containing core TF determinants of the neural crest lineage (*ZIC1*, *ZIC4*, *PAX3*). Lastly, NC.3 cells expressed a series of gene sets containing successively more mature markers of the neural crest lineage (e.g., *SOX9*, *NFIA*, *SOX10*, *TFAP2B*, *NRP1*, *GFRA3*).

In conclusion, transcriptome wide scRNA-seq of 7F-NMPs, presomitic mesoderm, dorsal neural tube, ventral neural tube and neural crest corroborates the high differentiation efficacy of each of our purposefully controlled downstream lineage protocols, notwithstanding the emergence of off-target neural crest in the dorsal neural tube protocol. All lineages expressed requisite markers for different stages of their specification, as well as post-cranial HOX genes (Extended Data Fig. 3d) including the thoracic HOX genes *HOXA9*, *HOXB9* and *HOXC9* (Extended Data Fig. 3e).

### Construction of 2.5D Posterior Axial Assembloids

Investigation of cell autonomous vs non-autonomous processes in complex tissues often deploy strategies such as engineered genetic mosaicism or cell-type specific promoters. 2.5D assembloid approaches, where different cell types are generated separately as 2D monolayers and subsequently assembled into complex 3D floating tissues^80^, allows these strategies to be deployed differently in different assembloid sub-compartments. For example, neuromuscular tissues could be generated where the neural compartment harbors an optogenetic tool, while the muscle compartment harbors a disease allele. To grant this capability in post-cranial tissues, we generated 2.5D assembloids comprised of all four of our 7F-NMP daughter lineages: dorsal neural tube, ventral neural tube, neural crest, and presomitic mesoderm (Fig. 4a). We began by extending the differentiation protocol of each lineage for an additional window of 192-264hrs (Fig. 4b). Dorsal neural tube and ventral neural tube were cultured in combined DAPT and EC23, inhibiting Notch and stimulating retinoic acid signalling respectively, to push cells out of the cell cycle. Additionally, ventral neural tube exposure to SAG was extended through this window. Neural crest was cultured in EC23, to further promote retinoic acid-mediated maturation. Lastly, presomitic mesoderm was cultured in 3µM CHIR, FGF2 and BME. This additional window of WNT signalling was critical for maintaining the purity of TBX6^+^ cells. That is, if CHIR was omitted between 192-264hrs or applied after a 24hr delay, the off-target caudal growth zone marker HB9^81^ was detected through expression of the HB9::GFP transgene present in our hPSC line^52^ (Extended Data Fig. 4a’-a’’). In contrast, application of a full 192-264hr window of CFB reduced HB9::GFP+ positive cells to less than 1%, and resulted in >90% TBX6^+^ cells (Extended Data Fig. 4a’’’).

**Figure 4.**
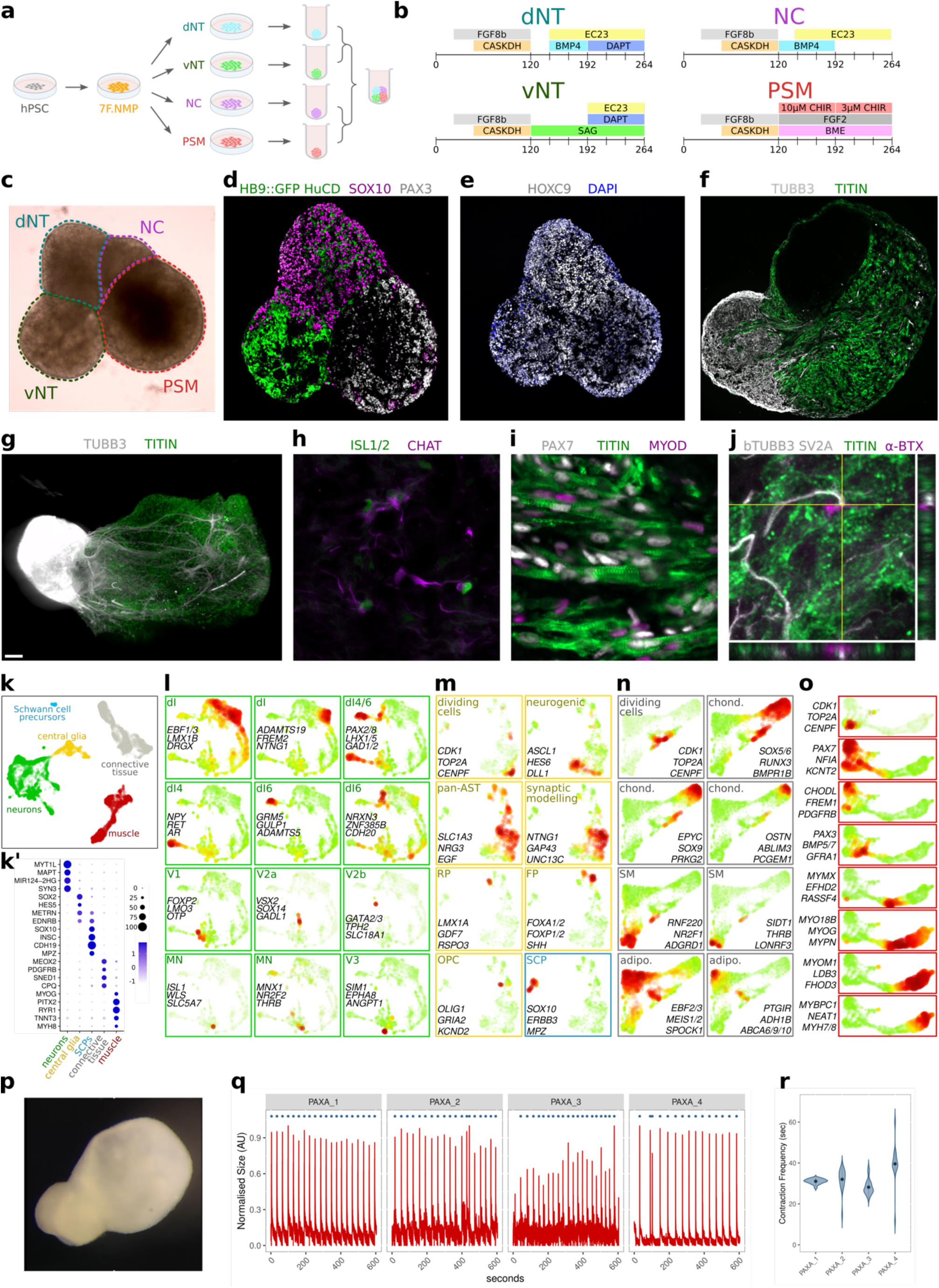
**(a)** Schematic of the 2.5D PAXA protocol. **(b)** Timing of factors used for the extended differentiation of the four 7F-NMP downstream lineages (dorsal neural tube, ventral neural tube, neural crest and presomitic mesoderm). **(c)** Brightfield image of a PAXA 24hrs after assembly (14DIV). Cryosections of a 15DIV PAXA immunostained for **(d)** neural (HuCD + endogenous HB9::GFP), neural crest (SOX10) and muscle progenitor (PAX3) markers, and **(e)** the posterior thoracic axial marker HOXC9. **(f)** 55DIV PAXA immunostaining shows spatial segregation of the neuronal (TUBB3) and muscle (TITIN) compartments. **(h)** Light sheet microscopy displaying nerve fibre tracts (TUBB3) extending into the muscle compartment (TITIN). Also present are **(h)** motor neurons (ISL1/2, CHAT), **(i)** muscle progenitors (PAX7, MYOD) and striated muscle fibres (TITIN), as well as **(j)** α-bungarotoxin staining located at the end of nerve fibres (co-stained with SV2a, TUBB3) amongst muscle fibres (TITIN). **(k)** snRNA-seq UMAP and Infomap cluster analysis of 55DIV PAXAs identifies five broad cell classifications that **(k’)** express requisite markers for neurons, Schwann cell precursors, central glia, connective tissues, and muscle. Subclustering of **(l)** only neurons, **(m)** central glia and Schwann cell precursors, **(n)** connective tissue, and **(o)** muscle cells. **(p)** Wholemount image of a 55DIV PAXA extracted from a 10min live video recording. **(q)** Normalised area of four PAXAs recorded over a 10min period. Spikes in area correspond to contraction events. **(r)** Contractions occur every ∼30-40sec and are statistically similar in all PAXAs.

Next, 2D monolayer cultures of each protocol were dissociated, re-aggregated into 3D spheroids, and subsequently combined into multi-lineage 2.5D assembloids that we term ***<u>p</u>***osterior ***<u>ax</u>***ial ***<u>a</u>***ssembloids, or PAXAs. Brightfield imaging of 14 days in vitro (DIV) PAXAs clearly displayed the four spheroids adhering together (Fig. 4c). Confocal imaging of 15 DIV PAXAs identified neural (combined signal from 488-HuCD, and endogenous HB9::GFP expressed in motor neurons), neural crest (SOX10^+^), and somitic mesoderm (PAX3^+^) compartments (Fig. 4d). Further, motor neurons could be identified in the ventral neural tube compartment through expression of HB9::GFP and ISL1/2 (Extended Data Fig. 4b). All compartments expressed the thoracic HOX gene, HOXC9 (Fig. 4e), corroborating their post-cranial identity. Confocal imaging of 55 DIV PAXAs (Fig. 4f-j) identified neural (TUBB3) and skeletal muscle (TITIN) sub-compartments (Fig. 4f), while light sheet imaging of cleared PAXAs displayed axonal processes extending into the muscle compartment (Fig. 4g, Supplementary Video 1). We also found co-expression of the motor neuron markers ISL1/2 and CHAT (Fig. 4h), the dorsal interneuron marker PAX2 (Extended Data Fig. 4c), the glial markers GFAP and S100β (Extended Data Fig. 4d), the muscle progenitor markers PAX7 and MYOD (Fig. 4i, Extended Data Fig. 4e), and striated organisation of multinucleated TITIN filaments (Fig. 4i, Extended Data Fig. 4e). Lastly, α-bungarotoxin, which binds to acetylcholine receptors clustered at the neuromuscular junction (NMJ), was detected abutting neural filaments co-stained for β-tubulin and SV2A (Fig. 4j).

### snRNA-seq of PAXAs identifies extensive tissue complexity

To systematically identify all cell types present, we next processed whole PAXA tissue using single-nuclear RNA-seq. UMAP analysis and graph based Infomap clustering of 9240 nuclei broadly classified five different cell types based on expression of requisite markers; neurons, central glia, Schwann cell precursors, connective tissue, and skeletal muscle (Fig. 4k-k’). All five cell clusters were represented in technical sequencing replicates (Extended Data Fig. 4f). Sub-clustering of neurons (Fig. 4l) identified gene sets containing markers for dorsal horn neurons (*EBF1/3*, *LMX1B* and *DRGX*), dorsal interneuron subtypes 4/6 (*PAX2/8*, *LHX1/5*, *NPY*, *GRM5*, *NRXN3*), V1 interneurons (*FOXP2*, *LMO3*, *OTP*), V2a interneurons (*VSX2*, *SOX14*), V2b interneurons (*GATA2/3*, *TPH2*), two gene sets marking motor neurons (*ISL1*, *WLS*, *SLC5A7*, *MNX*, *NR2F2*), and V3 interneurons (*SIM1*, *EPHA8*). Sub-clustering of central glia and Schwann cell precursors together (Fig. 4m) identified gene sets containing markers for actively dividing cells (*CDK1*, *TOP2A*), neurogenic astrocytes (*ASCL1*, *HES6*), pan-astrocytic markers (*SLC1A3*, *NRG3*), genes involved in synaptic modelling (*NTNG1*, *GAP43*), roof plate cells (*LMX1A*, *GDF*), two groups of cells expressing pan-floor plate markers (*FOXA1*, *FOXP2*) and one group also expressing lateral floor plate markers (*SULF1*, *ARX*, *CMTM8*, Extended Data Fig. 4g). Markers of cilia were expressed in all roof plate and floor plate cells (*CFAP47/54/61*, *DYNC2H1*, *ARMC2*, Extended Data Fig. 4h). Gene sets were also found that identified oligodendrocyte progenitor cells (*OLIG1*, *GRIA2*), and Schwann cell precursors (*SOX10*, *INSC*, *MPZ*) that appeared to be segregated into myelinating (*PMP22*) and non-myelinating (*EYA1*, *HAND2*, *FOLH1*) sub-types (Extended Data Fig. 4i-j). Sub-clustering of connective tissue (Fig. 4n) identified three different lineages, each of which contained a small number of actively dividing cells (*CDK1*, *TOP2A*). First, a chondrocyte lineage expressed gene sets containing critical transcription factor determinants (*SOX5/6/9*, *RUNX3*) and a gene set containing osteoblast markers (*OSTN*, *PCGEM1*). A second lineage expressed gene sets containing markers of smooth muscle (*RNF220*, *SIDT1*), while a third lineage expressed gene sets containing markers of brown adipocytes progenitors (*EBF1/2*, *MEOX1/2*, *SCARA5*, *ABCA6/9/10*), that developmentally originate from NMP derived dorsal myotome^82^. Lastly, sub-clustering of muscle cells (Fig. 4o) identified an axis of differentiating skeletal muscle. Located on the left of the UMAP were actively dividing cells (*CDK1*, *TOP2A*), as well as cells expressing gene sets marking muscle satellite cells (*PAX7*, *NFIA*), putative fibro-adipogenic progenitors (*CHODL*, *FREM1*, *PDGFRB*), proliferating satellite cells (*PAX3*, *BMP5/7*), fusion competent myoblasts (*MYMX*, *RASSF4*), pan skeletal muscle markers (*MYO18B*, *MYOG*, *MYPN*), developing contractile muscle fibres (*MYOM1*, *LDB3*, *FHOD3*), and mature muscle fibre markers (*MYBCP1*, *NEAT1*, *MYH7*/8)

### PAXAs display spontaneous contractions

Lastly, we recorded live videos of 55 DIV PAXAs for 10min periods (Fig. 4p-r). Computational analysis of PAXA size over time identified peaks that corresponded with spontaneous contractions (Supplementary Video 2) in all recorded PAXAs (Fig. 4q). The average contraction frequency was approximately 35sec and did not significantly differ between PAXAs (Fig. 4r). In summary, PAXAs constitute a complex in vitro tissue containing an extensive array of mature cell types from the dorsal neural tube, ventral neural tube, neural crest, and presomitic mesoderm lineages, while also displaying spontaneous, rhythmic and persistent functional contractions. Thus, through purposeful 2D differentiation of all NMP downstream lineages, followed by assembly and long-term 3D floating culture, PAXAs constitute a unique and tractable model of post-cranial tissue.

## DISCUSSION

We describe the deployment of 7F-NMPs and four downstream lineage protocols to construct human PAXAs: an advanced, multi-compartment, heterogeneous, posterior axial assembloid. The cell diversity seen in PAXAs has never been reported from any single NMP protocol and demonstrates the impressive multi-potency of 7F-NMPs. Further, the complexity of PAXA tissue lends itself to studying a wide range of biological processes unfolding in human posterior tissues. For example, but not limited to, post-mitotic maturation of neural subtypes in the spinal cord, synaptic modelling by central astrocytes, axonal transport, central/peripheral myelination, nascent NMJ formation, satellite cell homeostasis, the biology of muscle cell fusion, or mature muscle fibre metabolism. Moreover, since PAXAs are derived from tractable hPSCs in vitro, each of these processes could be investigated in different opto/chemo-genetically modulated states of neuromuscular activity, or different states of genetically/environmentally imparted pathology. We anticipate that PAXAs will have particular use in modelling diseases where complex pathologies unfold across multiple cellular compartments, such as motor neuron diseases, muscular dystrophies and sarcopenia. To facilitate uptake and exploration of our 7F-NMP and PAXA protocols towards these goals, we provide a browsable shiny app of all presented bulk-, single-cell-, and single-nuclear-RNA-seq datasets (https://nigelkee.shinyapps.io/PAXAs).

Embryonic development uses a small number of signals (e.g., WNT, FGF, BMP, sonic hedgehog, retinoic acid) in different permutations across space and time to specify different lineages. However, 3D organoid protocols use a single culture cocktail, at a given moment in time, to drive the specification of all emerging lineages. By contrast, our 2.5D assembloid protocol combines independently generated monolayer differentiations, allowing flexibility to specify cells unconstrained by the developmental requirements of neighbouring lineages. For example, temporally controlled BMP application can be used to specify both neural crest and dorsal neural tube, while dorso-ventral patterning in the neural tube and somite can be deployed independently. Further, genetically mosaic PAXAs would provide additional avenues to target cell types of interest. For instance, ventral neural tube neurons and their target muscle cells can be generated using different hPSC lines. Alternatively, imperfect cell-type specific promoters that drive expression of molecular tools can be introduced in one lineage without risking mis-expression in another. Further, purposeful construction using all or some of each lineage would ensure the presence of the desired lineages in every PAXA. That 7F-NMPs can derive all downstream lineages also greatly simplifies the engineering of human posterior tissues. Our protocols use the same coatings and base media, while long-term maturation of PAXAs does not require supplementation with additional growth factors, likely due to the production of multiple growth factors within PAXAs themselves, including *IGF1*, *HGF*, *AR*, *FGFs* and *NGFs* (Extended Data Fig. 5b).

Maintenance of stable 7F-NMPs allows for the timed triggering of downstream differentiation. In the current protocols, where differentiation is triggered at 120hrs, daughter cells express matched thoracic level HOX9 genes. Matched *HOX* levels between motor neurons and muscle have functional consequences, since transplantation experiments show that while unmatched motor neurons-muscle can establish initial connectivity, over time these NMJs degenerate^83^. Thus, efforts studying human NMJ pathology would ideally use in vitro tissues of a matched *HOX* code. Further, since co-linear *HOX* activation is occurring in 7F/6F-NMPs, triggering differentiation earlier or later than 120hrs would be expected to specify downstream lineages of different A-P identity. We achieve this result in preliminary experiments specifying ventral neural tube (Extended Data Fig. 6), including 6F dependant expression of HOXC10 (Extended Data Fig. 6d-e), further underscoring the authenticity of our protocols. Of note, multiple reports implicate extensive crosstalk between WNT and FGF^37^, and Notch^32^ in NMPs, and where *HOX* progression depends on the duration that a cell resides in the niche^15,16^ and the level of FGF signalling within the niche^84,19,85^. While we use constant concentrations of CHIR, FGF8b, and DAPT, the production of many effectors of these pathways by 7F-NMPs themselves confounds a mechanistic interpretation (Extended Dat Fig. 5a). Therefore, we anticipate that our 7F-NMP and four downstream lineage protocols will provide a tractable, scalable source of human material to enable such mechanistic studies, allowing molecular analysis of HOX regulation both within NMPs, and in their maturing downstream lineages.

Combined WNT and FGF signalling are implicated in multiple aspects of NMP biology including post-cranial specification through induction of CDX1/2^5^, and downstream specification of presomitic mesoderm through induction of a TBXT driven transcriptional program^61,86,34,87^ and promotion of EMT^88^. Given this overlap, how does the full 7F cocktail specify NMPs without triggering off-target presomitic mesoderm? The remaining factors (A 83-01, K02288, SIS3, DAPT, heparin) invite several speculations. Firstly, SMAD inhibition can suppress both TBXT levels^28^ and EMT^89^. Thus, strong SMAD inhibition at both extracellular (AK) and intracellular (S) points of the pathway may result in 4.5µM CHIR inducing only low levels of TBXT that are insufficient to drive EMT or specify presomitic mesoderm, but that still permit CDX dependent post-cranial specification. Secondly, DAPT also suppresses levels of TBXT^32^ and may similarly inhibit EMT and presomitic mesoderm induction without compromising post-cranial specification. Further, heparin has an established role in stabilising factors in tissue culture media^90,91^, and likely stabilises WNTs and FGFs produced by NMPs themselves (Extended Dat Fig. 5a), and FGF8b supplied in the 7F cocktail. FGF receptors can be differentially activated by FGF8b or FGF2^92^, which also have different requirements for heparin^93^. Since substitution of FGF8b for FGF2 increased TBX6^+^ cells in NMP induction (Extended Data Fig. 1e), and substitution of FGF2 for FGF8b decreased TBX6^+^ cells in presomitic mesoderm induction (Extended Data Fig. 2b’), FGF8b appears to be the superior ligand for 7F-NMP specification.

Noteably, some cell types were not well represented in our data, such as sensory neurons of the dorsal root ganglia, mature Schwann cell precursor derivatives, or myelinating oligodendrocytes. These cell types likely require additional modification of the early monolayer protocols, and/or longer-term floating culture to allow further cell maturation. Moreover, off-target neural crest comprised a large proportion of cells in the dorsal neural tube protocol, but this may be ameliorated by further optimising the timing of BMP4/EC23 application, and/or by additionally incorporating alternative BMP ligands^94,43^ or WNT signalling^45^. Further, the precise lineage relationship between the mature cell types seen in PAXAs, and cells from each of the four monolayer protocols, remains unresolved. Nonetheless, 7F/6F-NMPs display key hallmarks of authentic NMPs: expression of requisite markers conserved across species; temporal co-linear *HOX* induction in-line with A-P specification ranging from cervical through to lumbar levels; and genuine multi-potency to purposely derive all downstream lineages in ways that recapitulate signalling paradigms observed in vivo. Together, our 7F-NMP and PAXA protocols provide a uniquely versatile platform for the study of early development of posterior tissues, and for the in vitro modelling of a wide range of human diseases using an advanced, multi-compartment, posterior hPSC-derived assembloid.

## Acknowledgements

We would like to thank all authors and members of the Hedlund lab for valuable scientific discussions, feedback on the manuscript, and assistance in manuscript submission. Confocal microscopy was performed at the Biomedicum Imaging Core Facility (BIC) and the Imaging Facility at Stockholm University (IFSU). We would also like to thank Chris Molenaar, facility manager at IFSU, and Steven Edwards from the Advanced Light Microscopy facility at SciLifeLab, for providing excellent technical expertise on imaging. We also wish to thank Sonja Gustafsson and the Single Cell Core Facility for Flemingsberg campus (SICOF), Karolinska Institutet, for invaluable help in processing the Smart-seq3 bulk RNA-seq and scRNA-seq data. snRNA-seq was performed at SciLifeLab, Stockholm. Computations and data handling were enabled by resources provided by the National Academic Infrastructure for Supercomputing in Sweden (NAISS) and the Swedish National Infrastructure for Computing (SNIC) at UPPMAX. Thanks also to the Carmen Birchmeier laboratory, for the gift of the OLIG3 antibody. This work was supported by grants from the Swedish Research Council (2020-01049) to E.H., from the Department of Biochemistry and Biophysics, Stockholm University to E.H., from Hjärnfonden to E.H., from Åhlén’s Foundation (Åhlén stiftelsen) to N.K. and M.L., and Karolinska Institute Research Funds to N.K. N.K was supported by postdoctoral fellowships from Hjärnfonden (2017-2019) and the Swedish Society for Medical Research (SSMF, 2019-2021).

## Author contributions

N.K. conceived, designed and conducted differentiations for 7F-NMP, dorsal neural tube, ventral neural tube, neural crest, and presomitic mesoderm lineages, 2.5D PAXA differentiations, monolayer and PAXA immunohistology, confocal imaging, computational automated imaging analyses, bulk RNA-seq sample preparation, bulk-, single cell-, and single nuclear-RNA-seq based bioinformatic analyses, wrote the R based Shiny application, and wrote the manuscript. M.L. and S.G. conducted PAXA differentiations, PAXA immunos, and snRNA-seq PAXA processing. C.P. conducted all RNAscope experiments. I.M. conducted infomatic processing of PAXA contraction data. S.B. assisted in hPSC differentiations and sc-RNAseq processing. All authors contributed valuable scientific discussions. E.H. supervised all aspects of the project, including writing of the manuscript.

---

**Extended Data Figure 1.**
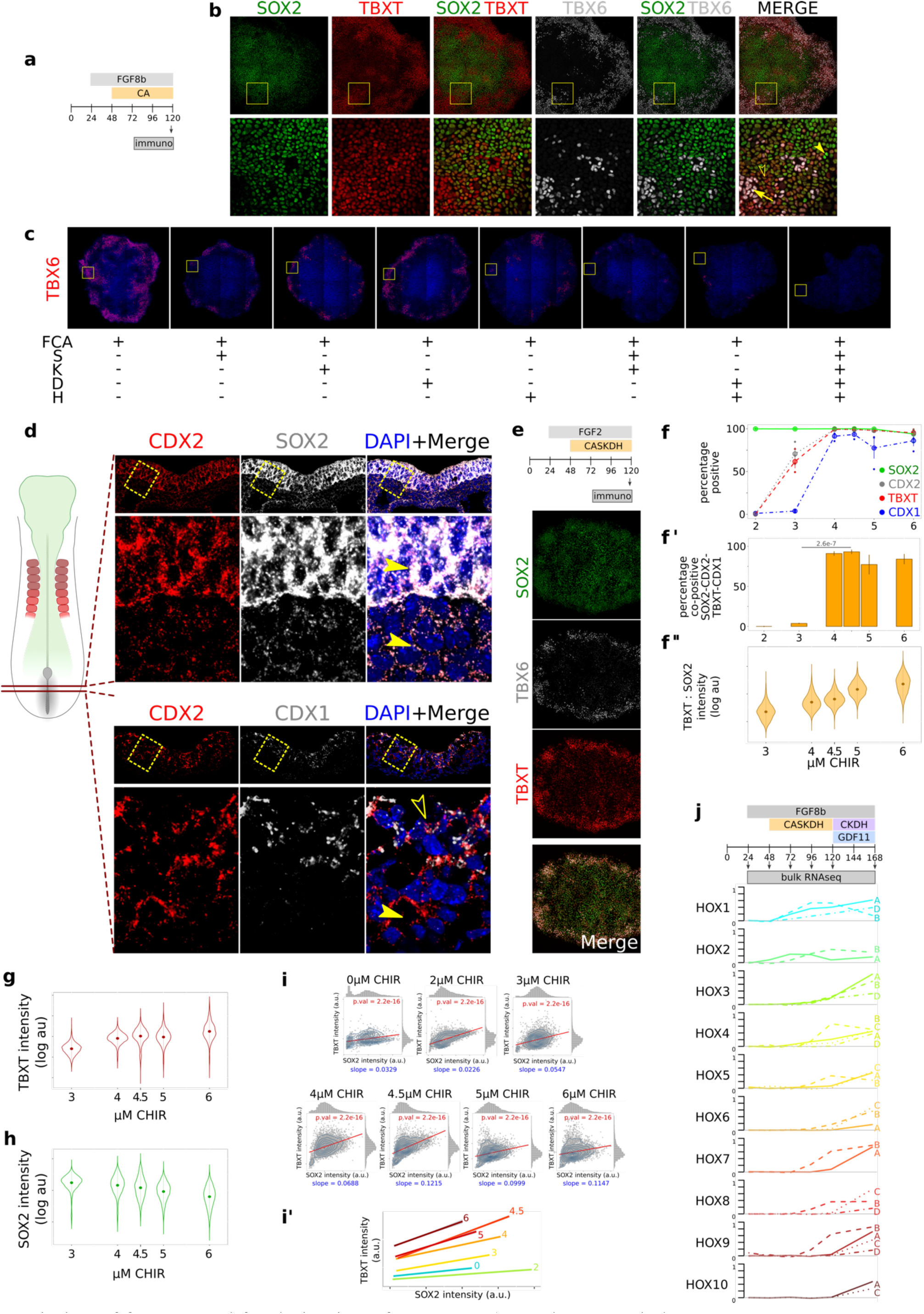
**a)** Timing of factors used for derivation of 3F NMPs (FGF8b at 50ng/ml, CHIR at 4.5µM, A 83-01 at 10µM). **b)** Immunostaining of the NMP markers SOX2 and TBXT, and the presomitic mesoderm marker TBX6. **c)** Representative confocal images of whole colonies that were analysed in Figure 1c-d. Boxed area shows the insert displayed in Figure 1c. **d)** RNAscope histology of mRNA probes targeting CDX2, SOX2 and CDX1 in E8.0 mouse caudal growth zone tissue. **e)** Equimolar substitution of FGF8b for FGF2 leads to emergence of off-target TBX6+ cells at the periphery of the colony. **f-f’’)** Behaviour of SOX2, CDX2, TBXT and CDX1 expression in an additional line, hiPSC 18a {Kiskinis:2014}, when differentiated in the 7F cocktail across a gradient of CHIR. These result phenocopy the original HB9::GFP hPSC line{Di Giorgio:2008}. **g)** Per-cell TBXT intensity up the CHIR gradient for the HB9::GFP hPSC line. **h)** Per-cell SOX2 intensity up the CHIR gradient for the HB9::GFP hPSC line. **i)** Per-cell plotting of TBXT vs SOX2 intensity up the CHIR gradient for the HB9::GFP hPSC line. Axes are relative and different between the polts. **i’)** Plotting of the linear models for all CHIR concentrations in i, on the same plot. **j)** Extension of the 7F-NMP cocktail to a second phase (6F) that includes the sacral TGFβ family member GDF11 and excludes the SMAD inhibitors A 83-01 and SIS3. Under these conditions the lumbar HOX genes HOXA10 and HOXC10 are specifically upregulated at 168hrs.

**Extended Data Figure 2.**
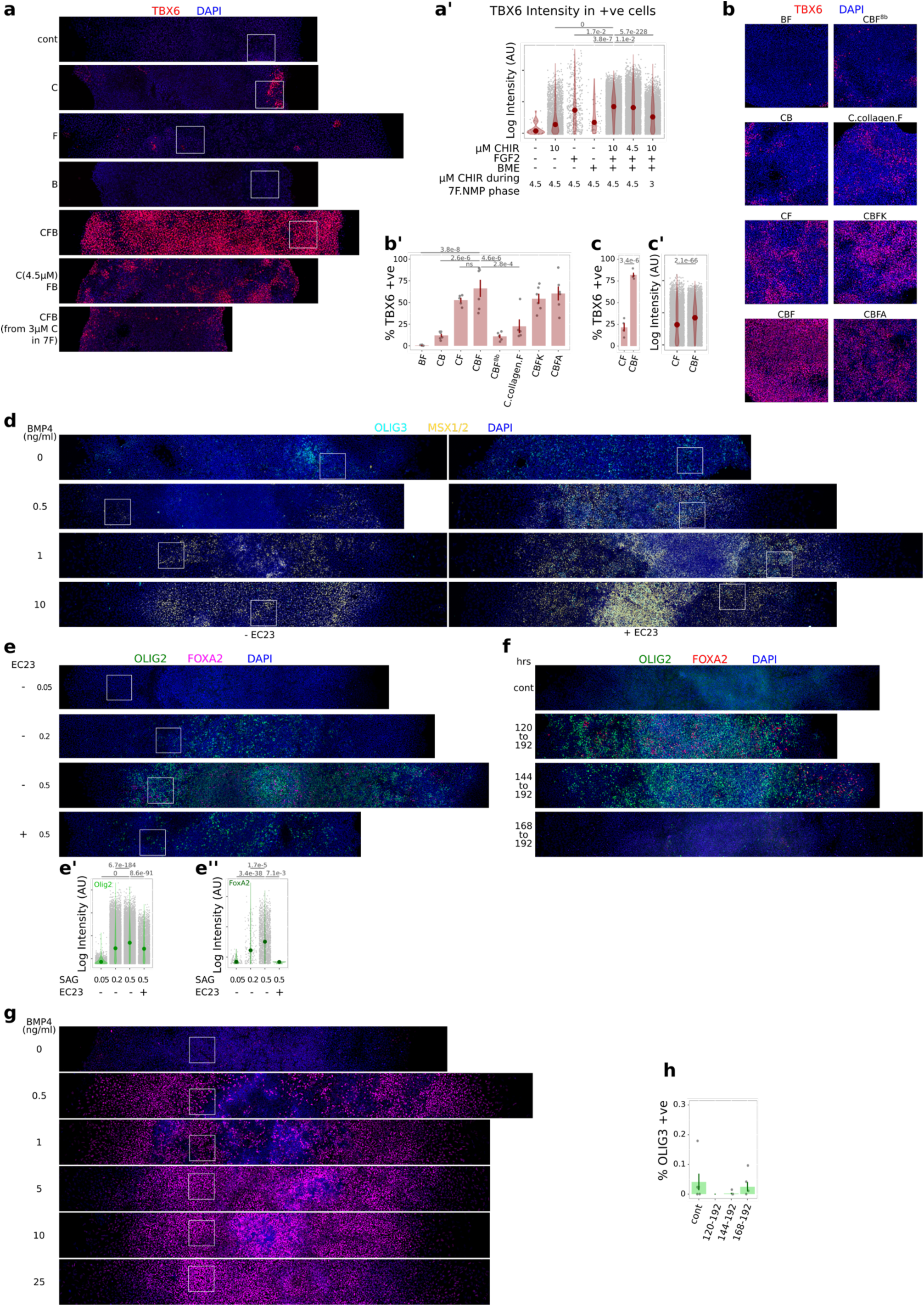
**a)** Full confocal images of representative TBX6 immunostainings from presomitic mesoderm permutations described in Figure 2b. White boxes denote the inserts pictured in the main figure. **a’)** Quantification of per cell TBX6 immunostaining intensities from Figure 2b. **b)** Confocal images of TBX6 immunostainings on an additional experiment testing presomitic mesoderm permutations. **b’)** Quantification of the percentage of TBX6+ cells from b. **c)** Quantification of the percentage of TBX6^+^ cells in repeat differentiations of presomitic mesoderm comparing CF to CBF. **c’)** Quantifications of per cell TBX6 immunostaining intensities from c. **d)** Full confocal images of representative OLIG3/MSX1/2 immunostainings on dorsal neural tube permutations from Figure 2c. **e)** Full confocal images of representative OLIG2/FOXA2 immunostainings on ventral neural tube permutations from Figure 2d. **e’)** Quantification of per cell OLIG2 immunostaining intensities from e. **e’’)** Quantification of per cell FOXA2 immunostaining intensities from e. **f)** Full confocal images of representative OLIG2/FOXA2 immunostainings on ventral neural tube permutations from Figure 2f. **g)** Full confocal images of representative SOX9 immunostainings on neural crest permutations from Figure 2g. **h)** Quantification of the percentage of off-target OLIG3^+^ cells when BMP4 is applied for different time windows after 7F-NMP exit. This OLIG3 Immunostaining was performed together with SOX9 in Figure 2h. C = CHIR, F = FGF2, B = basal matrix extract, F8b = FGF8b, collagen = rat Collagen 1, K = K02288, A = A 83-01.

**Extended Data Fig. 3.**
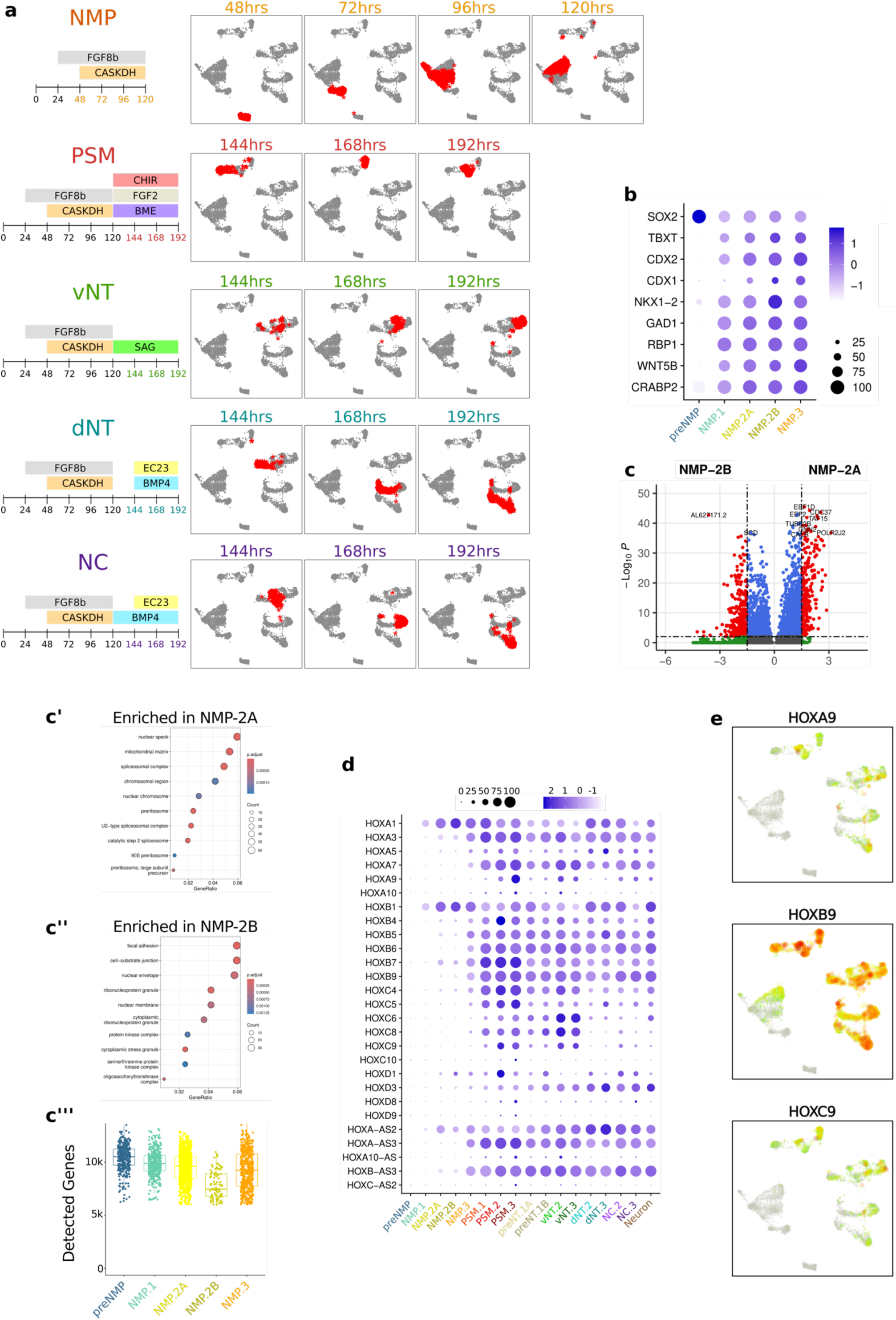
**a)** UMAP representations of the cells belonging to the samples taken from each protocol. Timepoints are indicated above each UMAP, with the cells from that timepoint marked with a red cross. **b)** Dot plot of requisite NMP markers shows robust expression in NMP1-3 cell clusters. **c’)** GOterms enriched in NMP.2A. **c’’)** GOterms enriched in NMP.2B. **c’’’)** NMP.2B cells have less detected genes than other NMP clusters, though we still detect ∼7000genes per cell. **d)** Expression of all detected HOX genes across all cell clusters. **e)** Expression of HOX9 genes. HOXB9 is robustly detected in 120hr 7F-NMPs, and in all later downstream lineages. In contrast, HOXA9 and HOXC9 are robustly detectable only in the downstream lineages.

**Extended Data Figure 4.**
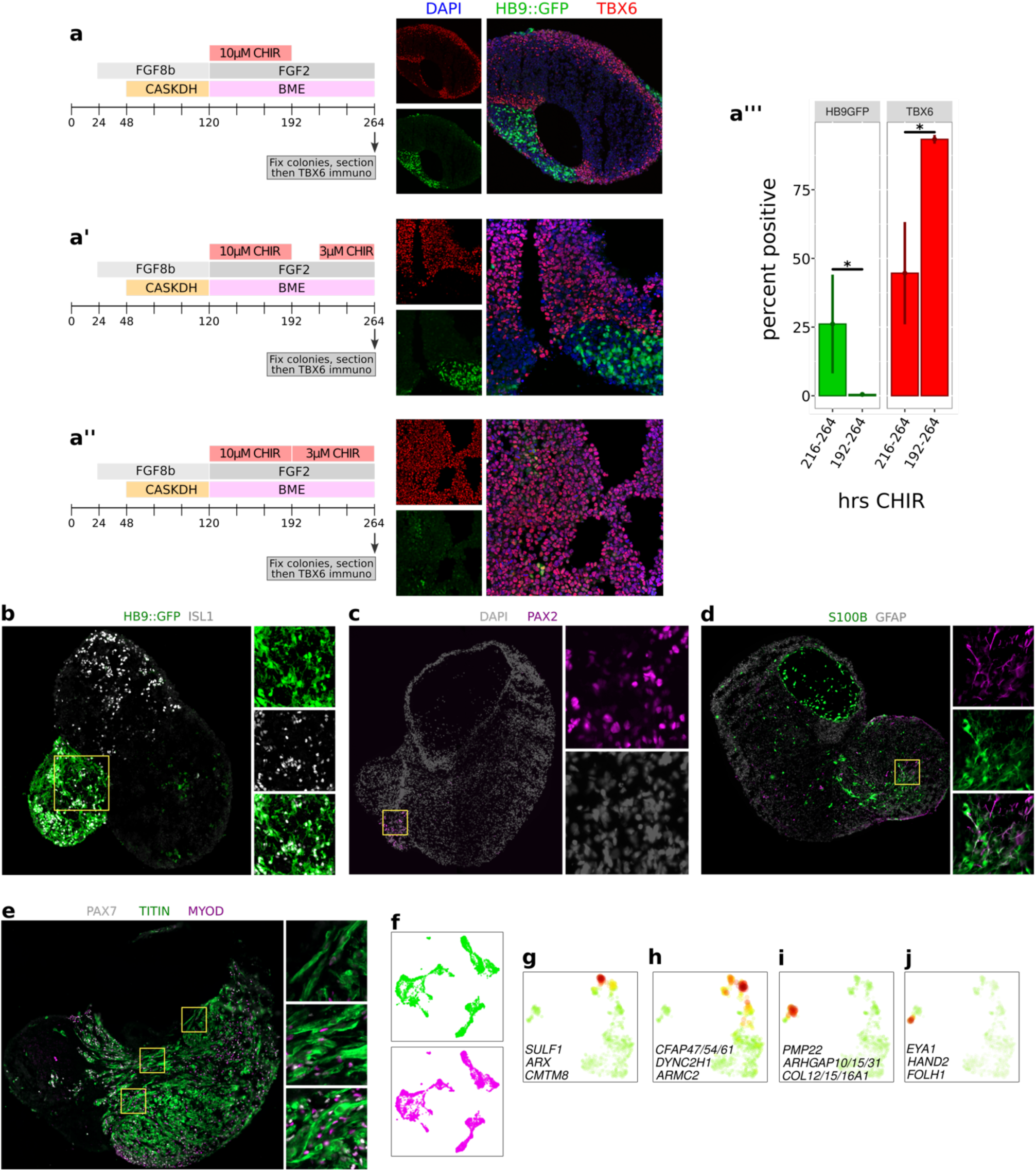
**a-a’’’)** LHS: Timing of factors used to extend the specification of presomitic mesoderm from 192-264hrs. MIDDLE: representative images of TBX6 stained cryostat sections of monolayer presomitic mesoderm colonies. RIGHT: quantification of decreased HB9::GFP expressing cells, and increased TBX6^+^ cells when applying CHIR for 192-264hrs (vs 216-264hrs). **b)** Immunostaining of 15DIV PAXA for ISL1 together with endogenous HB9::GFP signal shows double positive cells in the bottom left ventral neural tube compartment. **c)** Immunostaining of 55DIV PAXA for the dorsal neural tube interneuron marker PAX2. **d)** Immunostaining of 55DIV PAXA for the astrocytic markers S100β and GFAP. **e)** Immunostaining of 55DIV PAXA for the muscle progenitor markers PAX7 and MYOD, and the muscle fibre marker TITIN. Inserts show areas where the plane of sectioning illustrates multinucleated fibres displaying straited TITIN. **f)** snRNA-seq sequencing replicates (green and magenta) from independent lanes of a 10X genomics chip showing that cells from both lanes populate all cell clusters in the graph. **g)** A gene set containing putative lateral floor plate markers. **h)** A gene set containing cilia markers is expressed in all floor plate and roof plate cells. **i)** A gene set containing putative myelinating Schwann cell precursor markers. **j)** A gene set containing putative non-myelinating Schwann cell precursor markers.

**Extended Data Figure 5.**
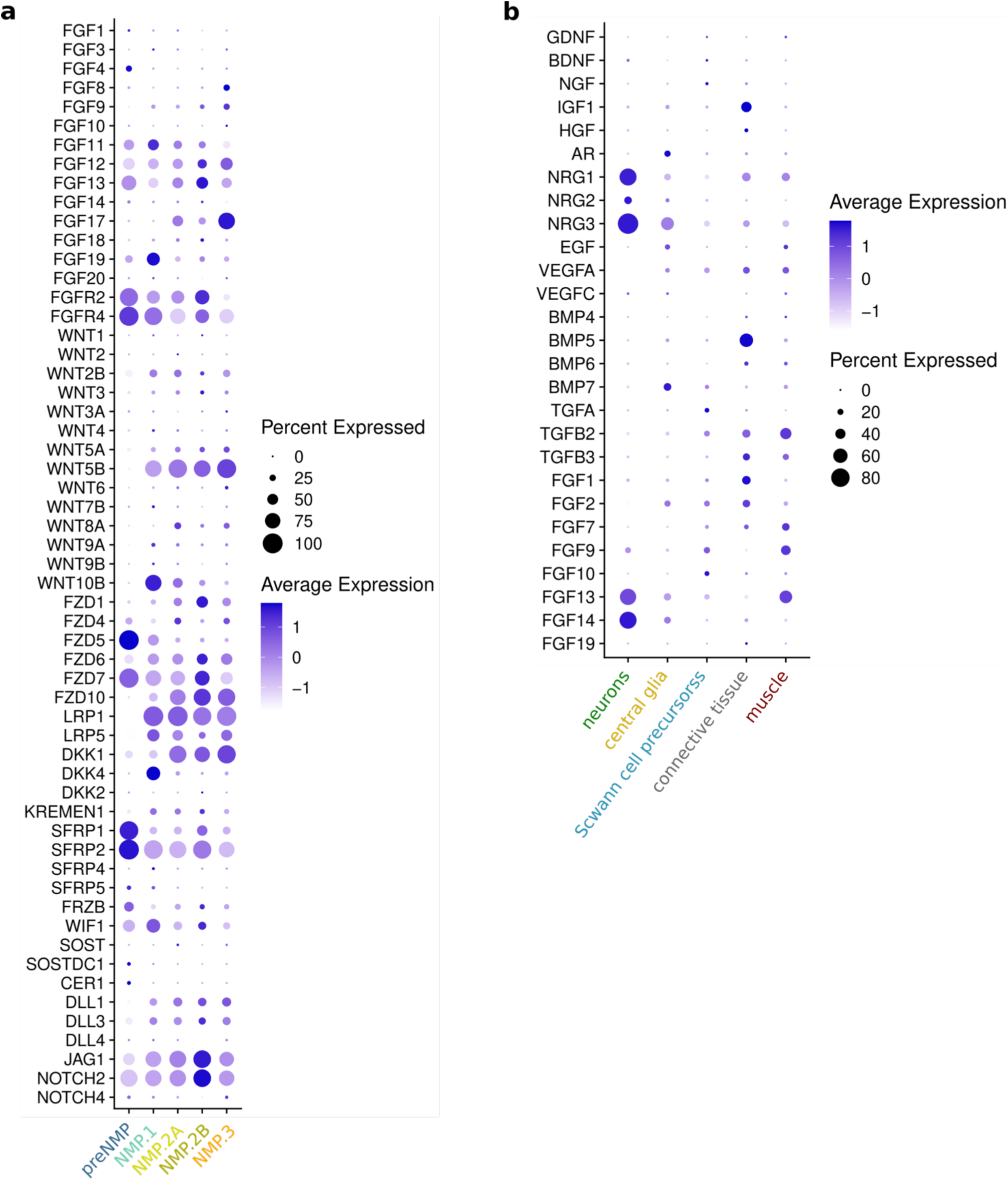
**a)** WNT, FGF and Notch pathway members expressed in monolayer 7F-NMP Smartseq3 scRNA-seq data. **b)** Growth factors expressed in 55DIV PAXA snRNA-seq data.

**Extended Data Figure 6.**
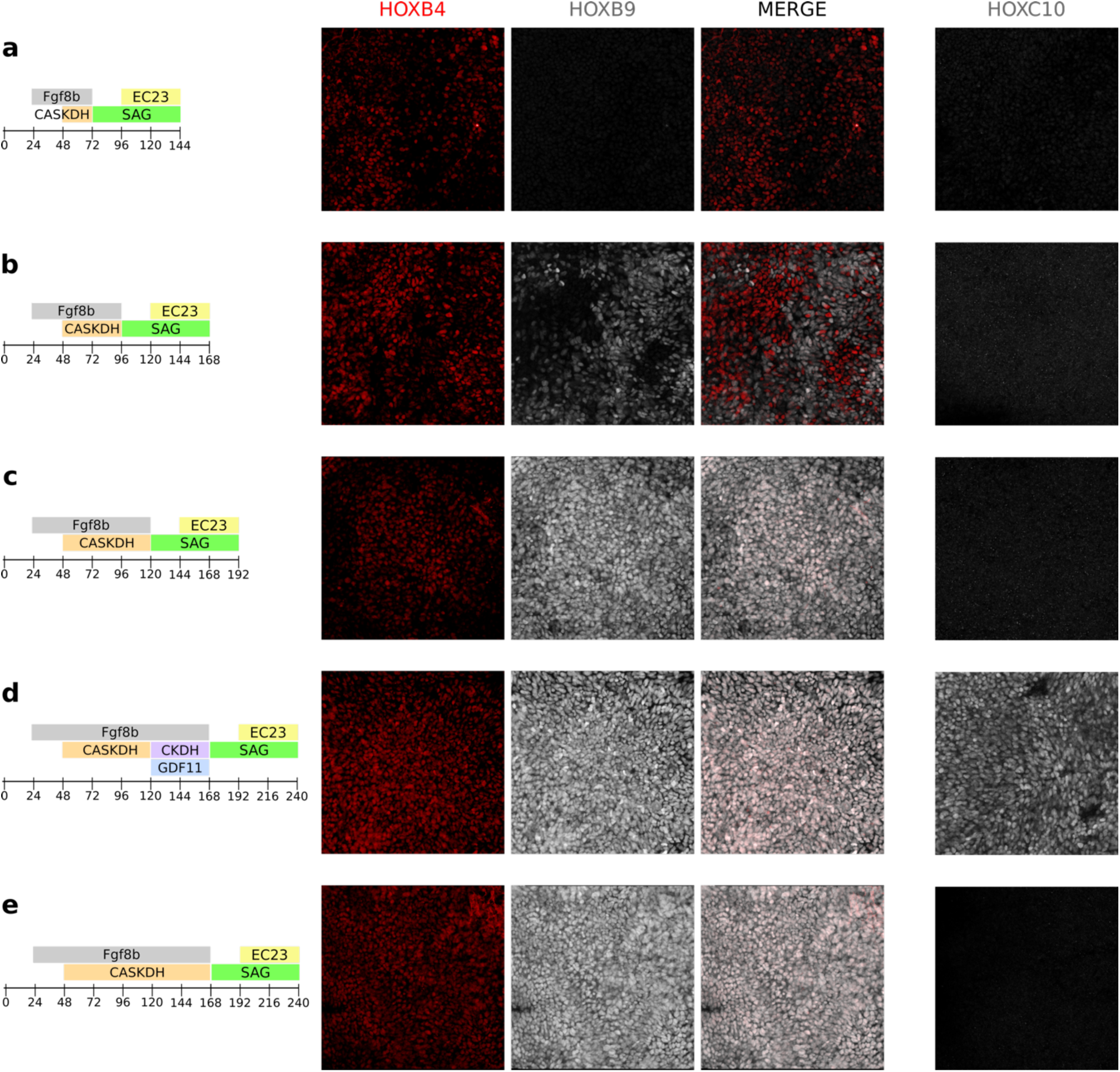
HOXB4, HOXB9, HOXC10 immunostaining in NMP derived ventral neural tube progenitors. Timing of factors used for each protocol depicted on the left. **(a)** HOXB4 staining is seen in ventral neural tube progenitors derived from all ages of NMPs. **(b)** In contrast, HOXB9 expression is seen in ventral neural tube progenitors beginning at 96hrs old 7F-NMPs, where it stains mostly HOXB4 negative cells. **(c)** By 120hrs old 7F-NMPs, all ventral neural tube progenitors are HOXB9+ but negative for HOXC10. **(d)** HOXC10 is first observed in 7F-to-6FNMPs (7F protocol followed by 48hrs of 6F), and **(e)** is not expressed in NMPs cultured for the same period in only 7F.

## References

1. Moris, N. et al. An in vitro model of early anteroposterior organization during human development. Nature 582, 410–415 (2020).

2. Olmsted, Z. T. & Paluh, J. L. Co-development of central and peripheral neurons with trunk mesendoderm in human elongating multi-lineage organized gastruloids. Nat. Commun. 12, 3020 (2021).

3. Pedroza, M. et al. Self-patterning of human stem cells into post-implantation lineages. Nature 622, 574–583 (2023).

4. Liu, L. et al. Modeling post-implantation stages of human development into early organogenesis with stem-cell-derived peri-gastruloids. Cell 186, 3776–3792.e16 (2023).

5. Wymeersch, F. J., Wilson, V. & Tsakiridis, A. Understanding axial progenitor biology in vivo and in vitro. Development 148, dev180612 (2021).

6. Solovieva, T., Wilson, V. & Stern, C. D. A niche for axial stem cells - A cellular perspective in amniotes. Dev. Biol. 490, 13–21 (2022).

7. Metzis, V. et al. Nervous System Regionalization Entails Axial Allocation before Neural Differentiation. Cell 175, 1105–1118.e17 (2018).

8. Hinoi, T. et al. Mouse Model of Colonic Adenoma-Carcinoma Progression Based on Somatic Apc Inactivation. Cancer Res 67, 9721–9730 (2007).

9. Franklin, V. et al. Regionalisation of the endoderm progenitors and morphogenesis of the gut portals of the mouse embryo. Mech. Dev. 125, 587–600 (2008).

10. Rothstein, M., Bhattacharya, D. & Simoes-Costa, M. The molecular basis of neural crest axial identity. Dev. Biol. 444, S170–S180 (2018).

11. Kinder, S. J. et al. The orderly allocation of mesodermal cells to the extraembryonic structures and the anteroposterior axis during gastrulation of the mouse embryo. Development 126, 4691– 4701 (1999).

12. Wymeersch, F. J. et al. Position-dependent plasticity of distinct progenitor types in the primitive streak. eLife 5, e10042 (2016).

13. Guillot, C., Djeffal, Y., Michaut, A., Rabe, B. & Pourquié, O. Dynamics of primitive streak regression controls the fate of neuromesodermal progenitors in the chicken embryo. Elife 10, e64819 (2021).

14. Tzouanacou, E., Wegener, A., Wymeersch, F. J., Wilson, V. & Nicolas, J.-F. Redefining the progression of lineage segregations during mammalian embryogenesis by clonal analysis. Developmental cell 17, 365–376 (2009).

15. Duboule, D. Temporal colinearity and the phylotypic progression: a basis for the stability of a vertebrate Bauplan and the evolution of morphologies through heterochrony. Development 1994, 135–142 (1994).

16. Lippmann, E. S. et al. Deterministic HOX patterning in human pluripotent stem cell-derived neuroectoderm. Stem cell reports 4, 632–644 (2015).

17. Ogura, T., Sakaguchi, H., Miyamoto, S. & Takahashi, J. Three-dimensional induction of dorsal, intermediate and ventral spinal cord tissues from human pluripotent stem cells. *Dev. (Camb.*, Engl.) 145, dev162214 (2018).

18. Libby, A. R. G. et al. Axial elongation of caudalized human organoids mimics aspects of neural tube development. Development 148, (2021).

19. Mouilleau, V. et al. Dynamic extrinsic pacing of the HOX clock in human axial progenitors controls motor neuron subtype specification. Development 148, dev194514 (2021).

20. Lee, J.-H. et al. Production of human spinal-cord organoids recapitulating neural-tube morphogenesis. *Nat*. Biomed. Eng. 6, 435–448 (2022).

21. Hackland, J. O. S. et al. Top-Down Inhibition of BMP Signaling Enables Robust Induction of hPSCs Into Neural Crest in Fully Defined, Xeno-free Conditions. Stem Cell Rep. 9, 1043–1052 (2017).

22. Frith, T. J. et al. Human axial progenitors generate trunk neural crest cells in vitro. Elife 7, e35786 (2018).

23. Fan, Y. et al. hPSC-derived sacral neural crest enables rescue in a severe model of Hirschsprung’s disease. Cell Stem Cell 30, 264–282.e9 (2023).

24. Gouti, M. et al. In Vitro Generation of Neuromesodermal Progenitors Reveals Distinct Roles for Wnt Signalling in the Specification of Spinal Cord and Paraxial Mesoderm Identity. PLoS Biol. 12, e1001937 (2014).

25. Gouti, M. et al. A Gene Regulatory Network Balances Neural and Mesoderm Specification during Vertebrate Trunk Development. Developmental cell 41, 243–261.e7 (2017).

26. Martins, J.-M. F. et al. Self-Organizing 3D Human Trunk Neuromuscular Organoids. Cell Stem Cell 26, 172–186.e6 (2020).

27. Pereira, J. D. et al. Human sensorimotor organoids derived from healthy and amyotrophic lateral sclerosis stem cells form neuromuscular junctions. Nat. Commun. 12, 4744 (2021).

28. Urzi, A. et al. Efficient generation of a self-organizing neuromuscular junction model from human pluripotent stem cells. Nat. Commun. 14, 8043 (2023).

29. Yaman, Y. I. & Ramanathan, S. Controlling human organoid symmetry breaking reveals signaling gradients drive segmentation clock waves. Cell 186, 513–527.e19 (2023).

30. Gribaudo, S. et al. Self-organizing models of human trunk organogenesis recapitulate spinal cord and spine co-morphogenesis. Nat. Biotechnol. 1–11 (2023) doi:10.1038/s41587-023-01956-9.

31. Gao, C. et al. Neuromuscular organoids model spinal neuromuscular pathologies in C9orf72 amyotrophic lateral sclerosis. Cell Rep. 43, 113892 (2024).

32. Cooper, F. et al. Notch signalling influences cell fate decisions and HOX gene induction in axial progenitors. Development 151, dev202098 (2024).

33. Akker, E. van den et al. Cdx1 and Cdx2 have overlapping functions in anteroposterior patterning and posterior axis elongation. *Dev. (Camb.*, Engl.) 129, 2181–93 (2002).

34. Koch, F. et al. Antagonistic Activities of Sox2 and Brachyury Control the Fate Choice of Neuro-Mesodermal Progenitors. Dev. Cell 42, 514–526.e7 (2017).

35. Blassberg, R. et al. Sox2 levels regulate the chromatin occupancy of WNT mediators in epiblast progenitors responsible for vertebrate body formation. Nat. Cell Biol. 24, 633–644 (2022).

36. Takemoto, T., Uchikawa, M., Kamachi, Y. & Kondoh, H. Convergence of Wnt and FGF signals in the genesis of posterior neural plate through activation of the Sox2 enhancer N-1. *Dev. (Camb.*, Engl.) 133, 297–306 (2005).

37. Amin, S. et al. Cdx and T Brachyury Co-activate Growth Signaling in the Embryonic Axial Progenitor Niche. Cell reports 17, 3165–3177 (2016).

38. Takemoto, T. et al. Tbx6-dependent Sox2 regulation determines neural or mesodermal fate in axial stem cells. Nature 470, 394–398 (2011).

39. Romanos, M. et al. Cell-to-cell heterogeneity in Sox2 and Bra expression guides progenitor motility and destiny. eLife 10, e66588 (2021).

40. Cunningham, T. J., Colas, A. & Duester, G. Early molecular events during retinoic acid induced differentiation of neuromesodermal progenitors. Biology open 5, 1821–1833 (2016).

41. Corral, R. D. del et al. Opposing FGF and Retinoid Pathways Control Ventral Neural Pattern, Neuronal Differentiation, and Segmentation during Body Axis Extension. Neuron 40, 65–79 (2003).

42. Zhao, X. & Duester, G. Effect of retinoic acid signaling on Wnt/β-catenin and FGF signaling during body axis extension. Gene Expr. Patterns 9, 430–435 (2009).

43. Iyer, N. R. et al. Modular derivation of diverse, regionally discrete human posterior CNS neurons enables discovery of transcriptomic patterns. Sci. Adv. 8, eabn7430 (2022).

44. Wind, M. et al. Defining the signalling determinants of a posterior ventral spinal cord identity in human neuromesodermal progenitor derivatives. Development 148, dev194415 (2021).

45. Gupta, S. et al. In vitro atlas of dorsal spinal interneurons reveals Wnt signaling as a critical regulator of progenitor expansion. Cell Rep. 40, 111119 (2022).

46. Chal, J. et al. Differentiation of pluripotent stem cells to muscle fiber to model Duchenne muscular dystrophy. Nat. Biotechnol. 33, 962–969 (2015).

47. Xi, H. et al. In Vivo Human Somitogenesis Guides Somite Development from hPSCs. Cell Rep. 18, 1573–1585 (2017).

48. Yamanaka, Y. et al. Reconstituting human somitogenesis in vitro. Nature 614, 509–520 (2023).

49. Diaz-Cuadros, M. et al. In vitro characterization of the human segmentation clock. Nature 580, 113–118 (2020).

50. Sáez, M. et al. Statistically derived geometrical landscapes capture principles of decisionmaking dynamics during cell fate transitions. Cell Syst. 13, 12–28.e3 (2022).

51. Anand, G. M. et al. Controlling organoid symmetry breaking uncovers an excitable system underlying human axial elongation. Cell 186, 497–512.e23 (2023).

52. Giorgio, F. P. D., Boulting, G. L., Bobrowicz, S. & Eggan, K. C. Human Embryonic Stem Cell-Derived Motor Neurons Are Sensitive to the Toxic Effect of Glial Cells Carrying an ALS-Causing Mutation. Stem Cell 3, 637–648 (2008).

53. Ikeya, M. & Takada, S. Wnt-3a is required for somite specification along the anteroposterior axis of the mouse embryo and for regulation of cdx-1 expression. Mech. Dev. 103, 27–33 (2001).

54. Kiskinis, E. et al. Pathways disrupted in human ALS motor neurons identified through genetic correction of mutant SOD1. Cell Stem Cell 14, 781–795 (2014).

55. Hagemann-Jensen, M. et al. Single-cell RNA counting at allele and isoform resolution using Smart-seq3. Nat Biotechnol 38, 708–714 (2020).

56. Hagey, D. W. & Muhr, J. Sox2 Acts in a Dose-Dependent Fashion to Regulate Proliferation of Cortical Progenitors. Cell Rep. 9, 1908–1920 (2014).

57. Kennedy, M. W. et al. Sp5 and Sp8 recruit β-catenin and Tcf1-Lef1 to select enhancers to activate Wnt target gene transcription. Proc. Natl. Acad. Sci. 113, 3545–3550 (2016).

58. Li, C.-M. et al. CTNNB1 Mutations and Overexpression of Wnt/β-Catenin Target Genes in WT1-Mutant Wilms’ Tumors. Am. J. Pathol. 165, 1943–1953 (2004).

59. Verrier, L., Davidson, L., Gierliński, M., Dady, A. & Storey, K. G. Neural differentiation, selection and transcriptomic profiling of human neuromesodermal progenitor-like cells in vitro. *Dev. (Camb.*, Engl.) 145, dev166215 (2018).

60. Albors, A. R., Halley, P. A. & Storey, K. G. Lineage tracing of axial progenitors using Nkx1- 2CreERT2 mice defines their trunk and tail contributions. Development 145, dev164319 (2018).

61. Liu, P. et al. Requirement for Wnt3 in vertebrate axis formation. Nat. Genet. 22, 361–365 (1999).

62. Neijts, R. et al. Polarized regulatory landscape and Wnt responsiveness underlie Hox activation in embryos. Gene Dev 30, 1937–1942 (2016).

63. Neijts, R., Simmini, S., Giuliani, F., Rooijen, C. van & Deschamps, J. Region-specific regulation of posterior axial elongation during vertebrate embryogenesis. Dev. Dyn. 243, 88–98 (2014).

64. Neijts, R., Amin, S., Rooijen, C. van & Deschamps, J. Cdx is crucial for the timing mechanism driving colinear Hox activation and defines a trunk segment in the Hox cluster topology. Dev. Biol. 422, 146–154 (2017).

65. Jurberg, A. D., Aires, R., Varela-Lasheras, I., Nóvoa, A. & Mallo, M. Switching Axial Progenitors from Producing Trunk to Tail Tissues in Vertebrate Embryos. Dev. Cell 25, 451–462 (2013).

66. McPherron, A. C., Lawler, A. M. & Lee, S.-J. Regulation of anterior/posterior patterning of the axial skeleton by growth/differentiation factor 11. Nat. Genet. 22, 260–264 (1999).

67. Veenvliet, J. V. et al. Mouse embryonic stem cells self-organize into trunk-like structures with neural tube and somites. Science 370, (2020).

68. Scott, L. E., Weinberg, S. H. & Lemmon, C. A. Mechanochemical Signaling of the Extracellular Matrix in Epithelial-Mesenchymal Transition. Front. Cell Dev. Biol. 7, 135 (2019).

69. Sasai, N., Kutejova, E. & Briscoe, J. Integration of Signals along Orthogonal Axes of the Vertebrate Neural Tube Controls Progenitor Competence and Increases Cell Diversity. PLoS Biol. 12, e1001907 (2014).

70. Lek, M. et al. A homeodomain feedback circuit underlies step-function interpretation of a Shh morphogen gradient during ventral neural patterning. *Development (Cambridge*, England) 137, 4051–4060 (2010).

71. Ribes, V. et al. Distinct Sonic Hedgehog signaling dynamics specify floor plate and ventral neuronal progenitors in the vertebrate neural tube. Genes Dev. 24, 1186–1200 (2010).

72. Liem, K. F., Tremml, G. & Jessell, T. M. A role for the roof plate and its resident TGFbetarelated proteins in neuronal patterning in the dorsal spinal cord. Cell 91, 127–38 (1997).

73. Müller, T. et al. The bHLH factor Olig3 coordinates the specification of dorsal neurons in the spinal cord. Genes Dev. 19, 733–743 (2005).

74. Marklund, U. et al. Detailed expression analysis of regulatory genes in the early developing human neural tube. Stem Cells and Development 23, 5–15 (2014).

75. Duval, N. et al. Msx1 and Msx2 act as essential activators of Atoh1 expression in the murine spinal cord. Development 141, 1726–1736 (2014).

76. Molotkova, N., Molotkov, A., Sirbu, I. O. & Duester, G. Requirement of mesodermal retinoic acid generated by Raldh2 for posterior neural transformation. Mech. Dev. 122, 145–155 (2005).

77. Wilson, L., Gale, E., Chambers, D. & Maden, M. Retinoic acid and the control of dorsoventral patterning in the avian spinal cord. Dev. Biol. 269, 433–446 (2004).

78. Martínez-Morales, P. L. et al. FGF and retinoic acid activity gradients control the timing of neural crest cell emigration in the trunk. The Journal of cell biology 194, 489–503 (2011).

79. Yang, J. et al. Guidelines and definitions for research on epithelial–mesenchymal transition. Nat. Rev. Mol. Cell Biol. 21, 341–352 (2020).

80. Pașca, S. P., et al. A nomenclature consensus for nervous system organoids and assembloids. Nature 609, 907–910 (2022).

81. Koronfel, L. M., Kanning, K. C., Alcos, A., Henderson, C. E. & Brownstone, R. M. Elimination of glutamatergic transmission from Hb9 interneurons does not impact treadmill locomotion. Sci. Rep. 11, 16008 (2021).

82. Rao, J. et al. Reconstructing human brown fat developmental trajectory in vitro. Dev. Cell 58, 2359–2375.e8 (2023).

83. O’Brien, M. K. & Oppenheim, R. W. Development and survival of thoracic motoneurons and hindlimb musculature following transplantation of the thoracic neural tube to the lumbar region in the chick embryo: Anatomical aspects. J. Neurobiol. 21, 313–340 (1990).

84. Liu, J.-P., Laufer, E. & Jessell, T. M. Assigning the Positional Identity of Spinal Motor Neurons Rostrocaudal Patterning of Hox-c Expression by FGFs, Gdf11, and Retinoids. *Neuron* **32**, 997– 1012 (2001).

85. Hackland, J. O. S. et al. FGF Modulates the Axial Identity of Trunk hPSC-Derived Neural Crest but Not the Cranial-Trunk Decision. Stem cell reports 12, 920–933 (2019).

86. Faial, T. et al. Brachyury and SMAD signalling collaboratively orchestrate distinct mesoderm and endoderm gene regulatory networks in differentiating human embryonic stem cells. *Dev. (Camb.*, Engl.) 142, 2121–2135 (2015).

87. Guibentif, C. et al. Diverse Routes toward Early Somites in the Mouse Embryo. Dev Cell (2020) doi:10.1016/j.devcel.2020.11.013.

88. Bulger, E. A., Muncie-Vasic, I., Libby, A. R. G., McDevitt, T. C. & Bruneau, B. G. TBXT dose sensitivity and the decoupling of nascent mesoderm specification from EMT progression in 2D human gastruloids. Development 151,.

89. Xu, J., Lamouille, S. & Derynck, R. TGF-β-induced epithelial to mesenchymal transition. Cell Res. 19, 156–172 (2009).

90. Furue, M. K. et al. Heparin promotes the growth of human embryonic stem cells in a defined serum-free medium. Proc. Natl. Acad. Sci. 105, 13409–13414 (2008).

91. Sasaki, N. et al. Heparan Sulfate Regulates Self-renewal and Pluripotency of Embryonic Stem Cells*. J. Biol. Chem. 283, 3594–3606 (2008).

92. Ornitz, D. M. & Itoh, N. The Fibroblast Growth Factor signaling pathway. Wiley Interdiscip. Rev. Dev. Biol. 4, 215–266 (2015).

93. Loo, B.-M. & Salmivirta, M. Heparin/Heparan Sulfate Domains in Binding and Signaling of Fibroblast Growth Factor 8b. J Biol Chem 277, 32616–32623 (2002).

94. Andrews, M. G. et al. BMPs direct sensory interneuron identity in the developing spinal cord using signal-specific not morphogenic activities. eLife 6, e30647 (2017).

